# Proteome plasticity in response to persistent environmental change

**DOI:** 10.1101/2021.03.22.436511

**Authors:** Matthew Domnauer, Fan Zheng, Liying Li, Yanxiao Zhang, Catherine E. Chang, Jay R. Unruh, Julie Conkright-Fincham, Scott McCroskey, Laurence Florens, Ying Zhang, Christopher Seidel, Benjamin Fong, Birgit Schilling, Rishi Sharma, Arvind Ramanathan, Kausik Si, Chuankai Zhou

**Author notes:** These authors contributed equally (Matthew Domnauer, Fan Zheng, Liying Li).

## Abstract

Temperature is a variable component of the environment and all organisms must deal with or adapt to temperature change. Acute temperature change activates cellular stress responses resulting in the refolding or removal of damaged proteins. However, how organisms adapt to long-term temperature change remains largely unexplored. Here, we report that budding yeast responds to long-term high temperature challenge by switching from chaperone induction to the reduction of temperature sensitive proteins and re-localizing a portion of its proteome. Surprisingly, we also find many proteins adopt an alternative conformation. Using Fet3p as an example, we find that the temperature-dependent conformational difference is accompanied by distinct thermostability, subcellular localization, and importantly, cellular functions. We postulate that in addition to the known mechanisms of adaptation, conformational plasticity allows some polypeptides to acquire new biophysical properties and functions when environmental change endures.

## Introduction

Temperature is an unstable parameter in the wild. Broadly, organisms deal with two types of temperature change: transient fluctuation or long-term temperature change that can persist for generations. Examples are, respectively, daily or seasonal shifts in temperature or a persistent rise in temperature because of climate change. Endothermic organisms maintain a constant body temperature. However, for the vast majority of ectothermic organisms, and some mammals, their body temperature co-fluctuates with the environmental temperature (Kisser and Goodwin, 2012). Temperature affects nearly all cellular processes. Proteins, the functional output of the genome, are metastable macromolecules and their folding, stability, and function are particularly sensitive to temperature. Therefore, how individual proteins, and the proteome at large, deal with temperature change remains a central question.

The cellular response to acute change in temperature has been studied extensively. For example, sudden increase in temperature, or heat shock, leads to large scale protein misfolding and aggregation (Hartl et al., 2011; Mogk et al., 2018; Ruan et al., 2017; Zhou et al., 2014). Cells have evolved mechanisms, including chaperones, proteasomes, and autophagy, to refold or remove the damaged proteins (Hartl et al., 2011). This energy-intensive process allows cells to survive acute temperature change by repairing or degrading the damaged proteome. In comparison, little is known about how cells adapt to extended shifts in temperature.

Some studies indicate that the adaptive strategies may be distinct between transient and persistent temperature shifts. For example, transcription of many heat shock proteins is initially induced upon temperature increase, but if the high temperature persists, the expression of heat shock proteins is reduced (Gasch et al., 2000). In fact, constant overexpression of HSF1, the master transcriptional regulator of the heat shock response, is detrimental to cells (Zheng et al., 2016). Similarly, chronic induction of chaperones in maladaptive stress response can lead to proteostasis defects and toxicity (Roth et al., 2014; Wang et al., 2006). Therefore, the adaptation to persistent environmental change is not merely an extension of the acute stress response program and cells likely rely on alternative strategies to cope with and thrive in a changed environment.

In addition to damaging existing proteins, temperature changes also affect *de novo* protein folding. Nascent polypeptides progress through an ensemble of kinetically and thermodynamically favorable intermediates of the natively folded structure (Onuchic et al., 1997). The energy landscape that defines the folding paths and final native structures of a protein is shaped by the physiochemical environment, speed of polypeptide synthesis, and availability of cofactors (Kramer et al., 2019; O’Brien et al., 2014). For example, synonymous mutations in the MDR1 gene, which change the decoding speed for the affected codons, result in a Mdr1 protein with different conformations and functions in cancer cells (Kimchi-Sarfaty et al., 2007). Likewise, co-translational folding of Fas2p *in vivo* requires the presence of its binding partner Fas1p (Shiber et al., 2018). Temperature change alters not only physiochemical parameters, but also the rate of translation and composition of the proteome, which together affect the *de novo* folding of proteins (Liu et al., 2013; Richter and Coller, 2015; Shalgi et al., 2013). These observations raise a fundamental question: how does the cellular proteome maintain both integrity and function upon a prolonged change in temperature?

## Results

### The proteome changes associated with adaptation to persistent high temperature

We set out to address this question using the budding yeast *Saccharomyces cerevisiae* for its adaptability and relatively small proteome. Yeast cells were cultured overnight at 21°C and then shifted to 35°C. The temperature change was initially associated with slow cell division (Figure 1A), protein aggregation (Figure 1B), and increased expression of some chaperones (Figure 1B and S1A). However, after 2-3 hours, the cells recuperated, and growth rate accelerated (Figure 1A). After 20hrs (∼15 generations) at 35°C, Hsp104-marked protein aggregates disappeared and many acute stress regulators, such as chaperones and trehalose synthases, returned to base line expression levels (Figure S1A, B).

**Figure 1:**
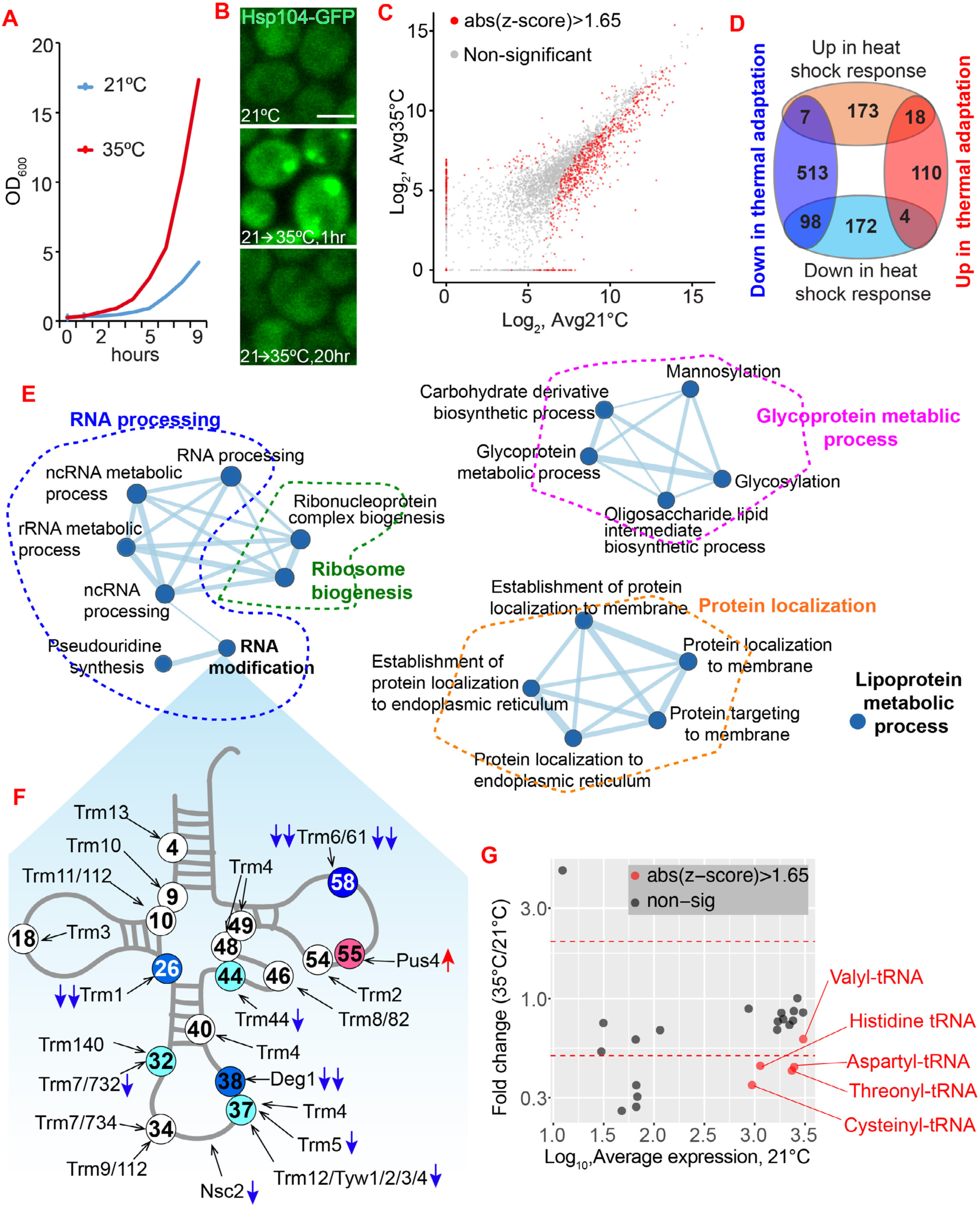
Adaptation to high temperature changes the expression of proteins. **A.** Representative growth curve of wild type cells at different temperatures. Cells were first grown at 21°C overnight and diluted and then transferred to 35°C. OD_600_ of the culture at various time points was calculated. **B.** Representative images of cells expressing Hsp104-GFP grown at different conditions. Images are representative of at least two independent experiments. Scale bar: 5μm. **C.** Expression level of individual proteins at both 21°C and 35°C from GFP screening. **D.** The overlap between proteins that respond to acute heat shock (15 min at 37°C, from (Gasch et al., 2000)) and thermal adaptation (20 hrs.). **E.** GSEA for the proteins down regulated during thermal adaptation depicted using Cytoscape (Shannon et al., 2003). The thickness of the connecting lines indicates the number of overlapping proteins between nodes. **F.** The expression changes of different tRNA modification enzymes. The tRNA and its modification sites (circled numbers) were compiled and modified from previous reviews (Hopper, 2013; Towns and Begley, 2012). Blue and red arrows indicate significantly decreased and increased expression, respectively. Double blue arrows indicate more than 2 folds reduction. **G.** The expression changes of different aminoacyl-tRNA synthetases. See also Figure S1.

To interrogate how the proteome adapts to the prolonged temperature increase without maintaining a canonical heat shock response, we performed a proteome-wide screen to track the expression and subcellular localization of individual proteins. We utilized a yeast GFP fusion library in which each strain has one of the 4,156 open reading frames tagged with GFP to report the protein’s abundance and localization (Huh et al., 2003). The GFP-tagged strains were grown either at 21°C or 35°C for 20hrs, followed by a high-content imaging screen (Figure S1C, D). The screen revealed significant differences in abundance of more than 700 proteins between the two temperatures (Figure 1C and Table S1). In addition to the reduction of chaperones, we found that many proteasome related proteins were reduced after 20hrs at 35°C (Figure S1E). Gene ontology analysis revealed attenuation of a broad spectrum of biological processes, including biogenesis of proteins and lipids (Figure 1E). Expression of some, but not all, tRNA modification enzymes and aminoacyl-tRNA synthetase was changed (Figure 1F, G), which may alter the aminoacyl-tRNA profile and decoding speed of certain codons. Among the lipid metabolism proteins, the ergosterol synthesis pathway was significantly reduced (Figure S1F). This is consistent with previous reports of temperature-dependent reduction of ergosterol, probably to maintain bulk membrane function at higher temperatures (Henderson et al., 2013; Parks and Casey, 1995). Most of these changes are not part of the acute heat shock response (Figure 1D) and are consistent with the general observation that more than 93% of gene expression changes associated with acute stress are not required to survive prolonged treatment with that stressor (Giaever et al., 2002; Yeger-Lotem et al., 2009).

### Thermal adaptation changes the subcellular localization of proteins

Subcellular localization is a determinant of protein function. Therefore, in addition to the changes in abundance, we also analyzed the subcellular distribution of proteins at 21°C and 35°C. Simple visual inspection revealed differential localization of many proteins between the two temperatures (Figure 2A, S2A). To quantitively evaluate protein localization changes, we trained a deep convolutional neural network that can recognize and classify patterns in digital images with high accuracy (DeepLoc) (Kraus et al., 2017; LeCun et al., 2015). Applying transfer learning we first optimized DeepLoc with a small subset of images of 15 subcellular compartments from the 21°C dataset (see Method for details). The transfer learning using our dataset was able to achieve high accuracy with an increasing number of training cells per localization class (Figure S2B), with several classes showing more than 95% accuracy (Figure S2C-E). t-SNE of the activations in the last convolutional layer of a testing dataset showed that the trained DeepLoc learned to distinguish different subcellular localizations and was able to arrange cells into separated populations (Figure 2B) (Maaten and Hinton, 2008).

**Figure 2:**
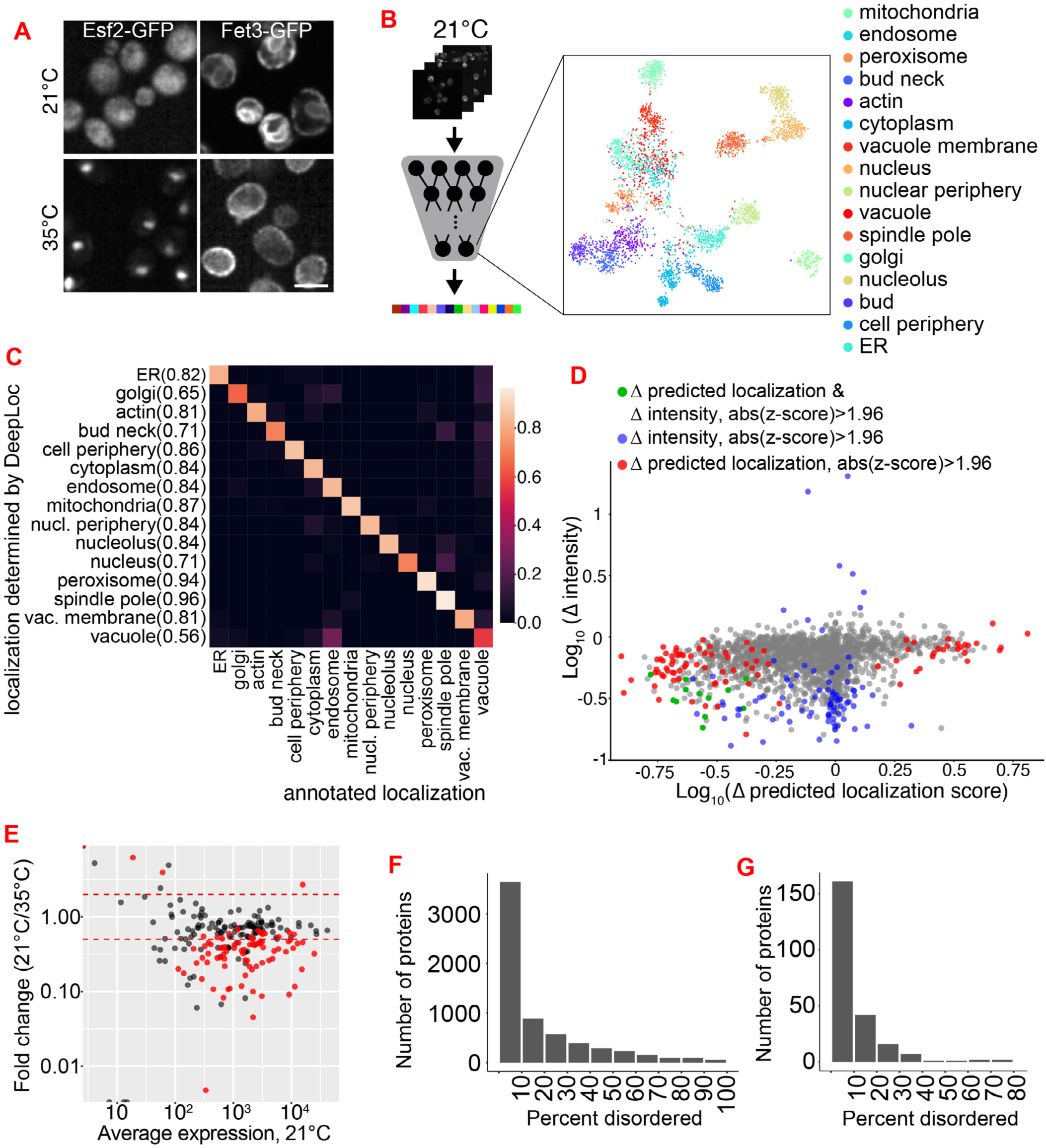
Subcellular localization changes of proteins and reduction of thermolabile proteins in response to persistent high temperature. **A.** Representative examples of altered subcellular localization. **B.** Schematic illustration of the transfer learning based on DeepLoc using dataset from 21°C samples and t-SNE plot of the activations in the last fully connected convolutional layer for 3866 single cells from the test set. **C.** Confusion Matrix of strains annotated to have a single localization (Huh et al., 2003). For each strain, the localization class that got the highest mean prediction score was set as the predicted localization of that strain. The prediction accuracy of trained DeepLoc is presented as the fraction of strains from each annotated class predicted to be certain class. Number in the brackets of y-axis is the recall value for each class. **D.** Localization and expression changes (Δ) for each strain between 21°C and 35°C. Each dot represents one strain. For each strain, the mean predicted localization score was calculated from single cells (N>20) for both 21°C and 35°C. The change in mean predicted localization score (35°C minus 21°C) for each strain was plotted on x-axis. The expression difference between two temperatures were normalized to expression value at 21°C (y-axis). **E.** Expression changes of aggregation-prone proteins during thermal adaptation. Red dots are proteins with absolute z-score larger than 1.65. Red dashed lines indicate the 2 folds increase or decrease. **F**, **G**. The percentage of residues in a given protein predicted to be intrinsically unstructured by IUPred2A (Mészáros et al., 2018). The distribution of proteome-wide (F) and the aggregation-prone proteins (G) were shown. There is no enrichment of intrinsically unstructured proteins among these aggregation-prone proteins. See also Figure S2.

We applied the trained DeepLoc to the entire dataset from both temperatures and compared the localization of proteins from the 21°C dataset with that of the manually curated localization database (Huh et al., 2003). DeepLoc predicted protein localizations with an average accuracy of >80% (Figure 2C). Comparison of protein localizations between 21°C and 35°C revealed that the cellular distribution patterns of ∼100 proteins significantly changed upon adaptation to high temperature (Figure 2D, Table S1). Some of these changes were likely due to the reduced expression at 35°C; however, for others, localization changes did not correspond with expression changes (Figure 2D). Taken together, these observations suggest many proteins change their subcellular distribution under persistent shift in temperature, either to perform new functions or to protect from thermal instability, or both.

### Reduction of aggregation-prone proteins

A key attribute of long-term temperature adaptation is the disappearance of protein aggregates. Previously we characterized protein aggregates induced by acute heat shock and identified 319 proteins inside the aggregates (Ruan et al., 2017). Out of these 319 aggregated proteins, 231 proteins were imaged in our current screen. Interestingly, expression of most of them was reduced after thermal adaptation with 99 of them showing more than a 2-fold reduction after thermal adaptation (Figure 2E). The aggregation-prone proteins are not enriched with intrinsically disordered proteins (Figure 2F, G, S2F). These results suggest that yeast cells adapt to the persistent high temperature by reducing the load of thermolabile proteins and relocating some proteins, which likely minimizes protein misfolding/unfolding at high temperature.

### Changes in protein conformation

Finally, we probed for changes in protein conformation since protein folding is sensitive to the molecular environment and temperature. As protein isolation may influence or enrich a specific protein conformation, we sought to assess protein conformation changes using limited proteolysis of the whole cell lysate in its native state. The accessibility of different residues by protease is an established method to probe protein conformation (Figure 3A) (Feng et al., 2014; Fontana et al., 1999, 2004). Three biological replicates of cell cultures grown for 20 hours at 21°C or 35°C were lysed via cryogrinder and clarified by high-speed centrifugation. The soluble fraction of lysate from both temperatures was briefly digested with proteinase K (PK) at room temperature and the cleaved peptides were identified using mass spectrometry (Figure 3A and see Method for details). Out of 2,364 proteins identified across six samples (Table S2), we focused on the 893 proteins that were identified in all samples (Table S2, Figure S3A). From the peptides, we deduced the PK cleavage sites and mapped these sites to the primary sequence of proteins to generate a 1D profile for each protein (Figure 3A). The shift between 1D profiles of the same protein from two conditions was used to estimate the change (differential score) in protein conformation (Figure S3B).

**Figure 3:**
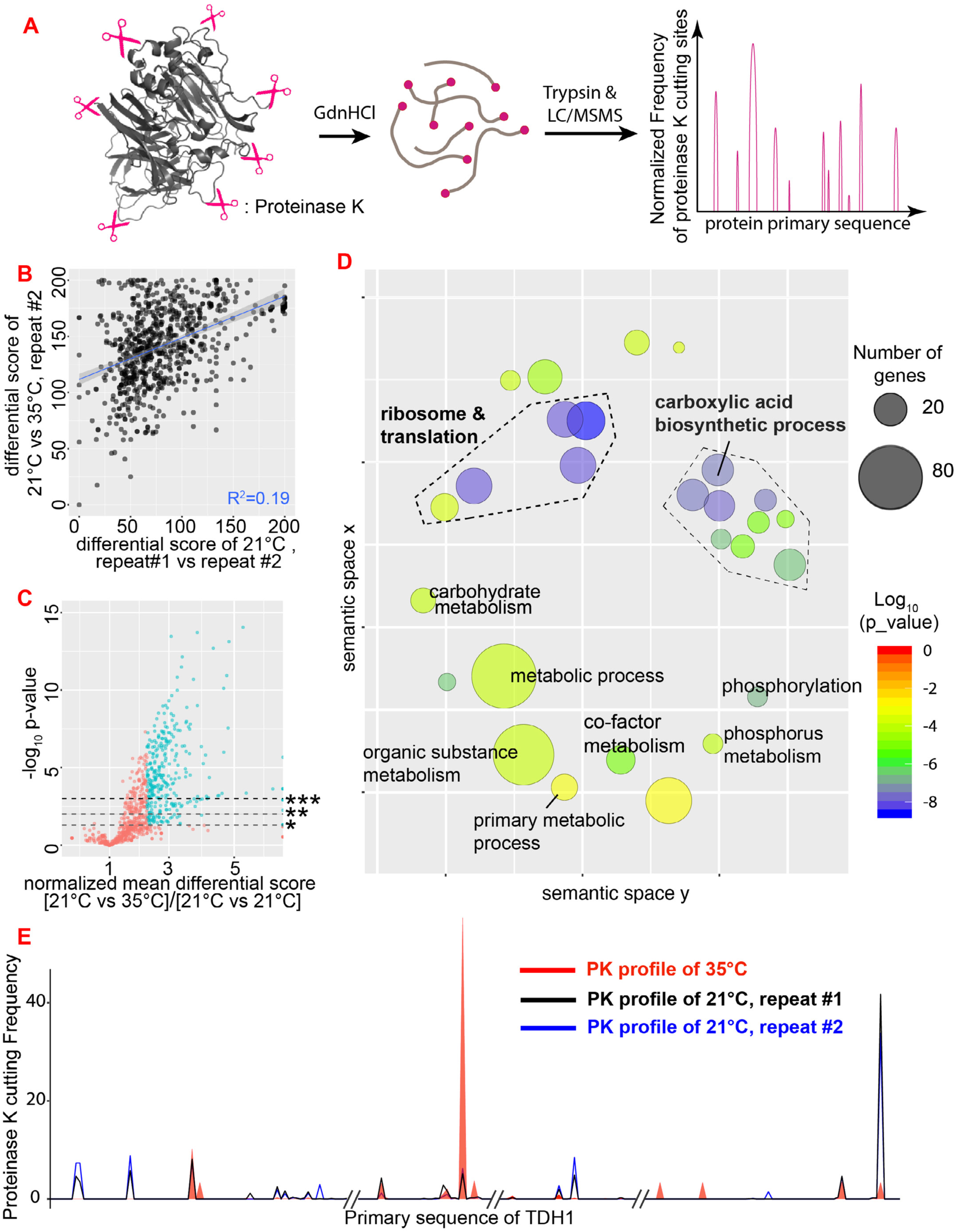
Limited proteolysis-coupled mass spectrometry reveals protein conformation changes. **A.** Schematic illustration of limited proteolysis method. Proteins in total cell lysate were briefly digested with proteinase K. The resulting peptides were identified by mass spectrometry and the proteinase cut sites were mapped to their primary sequence. **B.** The proteome-wide distribution of differential score between biological repeats of 21°C or between samples from 21°C and 35°C. Each dot represents a single protein. **C.** Volcano plot of temperature-driven differential score changes against p-value (ANOVA test). The temperature-driven differential score changes were normalized by the differential scores between biological repeats of 21°C. Horizontal gray lines denote p-value level of 0.05 (*), 0.01 (**), and 0.001(***). The proteins that showed at least two folds temperature-driven changes in differential scores were colored as cyan. **D.** GO term analysis of the cyan colored proteins in (C). The size of the bubbles indicates the number of proteins in each GO term and the bubble colors are coded according to its significance of enrichment. **E.** 1D profile of a protein that shows temperature-driven differential score change. The normalized PK-cutting frequency of each individual residues was plotted against the primary sequence of the protein. Two biological repeats from 21°C (blue and black traces) and one repeat from 35°C (red trace) were plotted together to illustrate the temperature-driven changes in multiple regions of the protein. The x axis is fragmented in order to visualize the peaks. See also Figure S3.

We first determined the variability in protease accessibility under the same experimental condition. The differential scores among biological repeats of 21°C samples centered around 70 (the range is 0-200; 0 means no difference and 200 means completely different), which sets the average noise of the assay (Figure 3B). There was no obvious correlation between abundance or intrinsic disorder of a protein and noise (Figure S3C, D) (Jones and Cozzetto, 2015). Approximately 75% of the detected proteins showed minimal difference in 1D profile between two temperatures, implying most proteins fold similarly as one would expect (Figure 3C, S3E, F). Surprisingly, for 228 proteins, we observed a considerable shift in 1D profile between two temperatures (Figure 3C, Table S3). Many of these proteins are metabolic enzymes and proteins involved in translation (Figure 3D). For example, Tdh1p and Sec13p have little variation between biological repeats of 21°C samples (Figure 3E, S3G, blue and black traces), while the proteins produced at 35°C show large shift of the 1D profile across their primary sequences (Figure 3E, S3G, red). This temperature-dependent 1D profile shift is not a function of protein abundance (Figure S3H, I) or their predicted melting point, one of the indicators of protein stability (Tm, Figure S3J) (Leuenberger et al., 2017).

### Alternative interaction vs alternative folding

The change in protease accessibility could be due to different intermolecular interactions, folding differences, or both (Figure 4A, B). To distinguish between these possibilities, we mapped the PK cleavage sites to the experimentally solved 3D reference structures of yeast proteins available from the PDB database and analyzed the depth of these PK sites relative to the surface of the protein (Figure 4C, 3D profiling, see Method for details). The alternative interactome predicts that PK sites from both temperatures should be restricted to the protein’s 3D surface, albeit to different regions (Figure 4A). On the other hand, the folding difference predicts that some of the internal residues at one temperature may be accessible to proteinase K at the other temperature (Figure 4B). By mapping the experimental PK sites to the 13,486 protein structures available for about 1,293 proteins (June, 2019), we observed three categories of proteins: (1) PK sites mapped to the surface of the reference structure at both temperatures (Figure 4D, G, H, and NP_011651 in movie S1)(Whitby et al., 2000); (2) PK sites from 21°C samples mapped to the surface, but were deep into the core of reference structure at 35°C (Figure 4E, S4A, and NP_014769 in movie S1) or *vice versa* (Figure 4F, S4B, and NP_011467 in movie S1); and (3) PK sites mapped to the core of the reference structure at both temperatures (Figure S4C and NP_013127 in movie S1).

**Figure 4:**
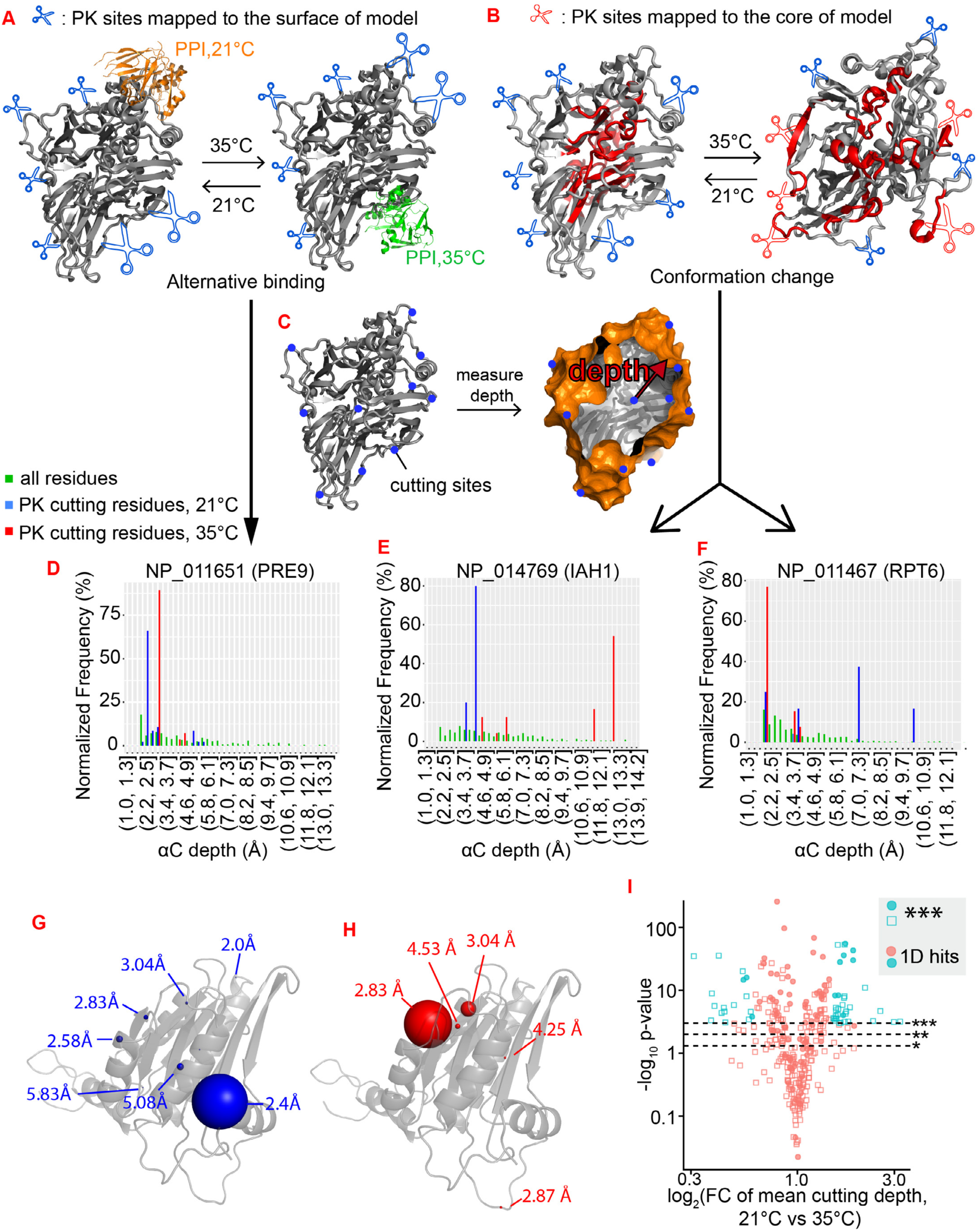
Identification of alternatively folded proteins. **A, B.** Schematic illustration of two alternative possibilities that can cause 1D profile shift. Alternative protein interactions (green and orange proteins in (**A)**) can shift the PK cutting sites along the surface of reference structure. The alternative conformation (**B**) can convert some inner residues (red) into PK-accessible residues. Illustration was modified from reference structure 1zpu (gray). **C.** Schematic illustration of mapping the PK sites relative to the surface of reference 3D structure of this protein to calculate the depth of each PK site (blue dots). Alpha carbon was used to represent the cutting depth of the residue. Illustration was modified from 1zpu (PDB) (Taylor et al., 2005). **D.** Representative example of a protein where PK sites were mapped to the surface of its reference structure at both temperatures. The distribution of alpha carbon depth of the entire structure is plotted in green as reference. The frequency of each PK site is normalized to the total number of cutting events in a given protein. See also movie S1 for the map of PK sites in the 3D structure. **E, F.** Example of proteins with differential PK sites at two temperatures. The distribution of alpha carbon depth of the entire structure is plotted in green as reference. See also movie S1 for the map of PK sites in the 3D structure. **G, H.** The localization and depth of PK sites in NP_011651 from 21°C (G) and 35°C (H) samples. The size of ball indicates the frequency of each PK site. Illustration was modified from 1fnt (PDB) (Whitby et al., 2000) **I.** Volcano plot of temperature-driven cutting depth changes against p value (ANOVA). Shown is the fold change of mean cutting depth of 21°C samples vs 35°C samples. Horizontal gray lines denote p value level of 0.05 (*), 0.01 (**), and 0.001(***). The proteins that show at least 1.5-fold changes were colored as cyan. The circles represent the proteins selected as hits in 1D profiling analysis. See also Figure S4.

We reasoned that category one likely represents the proteins with different interactomes between the two temperatures. Out of 228 candidates with a significant 1D profile shift, 96 had available 3D structures and were analyzed in 3D profiling, and 85 of them were in category one (Figure 4I). Thus, most of the observed 1D profile shifts likely originate from a change in protein-protein interactions (PPI) between two temperatures (Table S3). The PK-accessible sites in category three proteins were situated deep inside the protein based on the predicted structure (Figure S4C). Since most of these proteins are not predicted to be disordered and have a defined and reproducible 1D profile (Figure S4D, E), it is possible that the reference structures, determined under specific experimental conditions, do not capture the endogenous conformations (Dishman and Volkman, 2018; Tokuriki and Tawfik, 2009).

The proteins in category two are particularly interesting. At one of the temperatures, the PK sites were restricted to the surface of the reference structure, indicating that the reference structure captures the endogenous conformation of the protein at that temperature. For these proteins, the deep PK accessibility at the other temperature suggests a different folding state than the reference structure and that some of the internal residues in the reference structure are exposed on the protein surface at that temperature (Figure 4E, F). We selected 61 proteins that consistently showed differential 3D profiles upon temperature shift (Figure 4I, S4A, B, Table S4). We searched for nine different post-translational modifications (PTM), such as mono-, di-, tri-methylation, phosphorylation, ubiquitination, and acetylation. Out of 61 proteins, only 20 contained PTMs (Table S4). In addition, for each PTM-positive protein, only a small fraction of the total protein was PTM-modified (Table S4, Figure S4F). Therefore, most of the 3D profile shift cannot be accounted by a difference in PTMs.

To further evaluate possible conformational differences, we examined the thermostability of these proteins using the FastPP assay. The FastPP assay utilizes thermolysin, a hydrophobic residue-specific protease, to reveal the kinetics of thermal denaturation of proteins by degrading the denatured proteins (Minde et al., 2012) (Figure 5A). We found that many of these 61 proteins that showed a shift in their 3D profile also showed a reproducible difference in their thermostability (Figure 5B, S5A vs S5C, D). Moreover, in both limited proteolysis and FastPP assays, the protein samples were prepared at 4°C and then brought to room temperature (23°C) before adding proteases. Therefore, the observed 3D profile and thermostability differences indicate that the *in vivo* conformation adopted by these proteins is stable enough to withstand a temperature shift *in vitro*.

**Figure 5:**
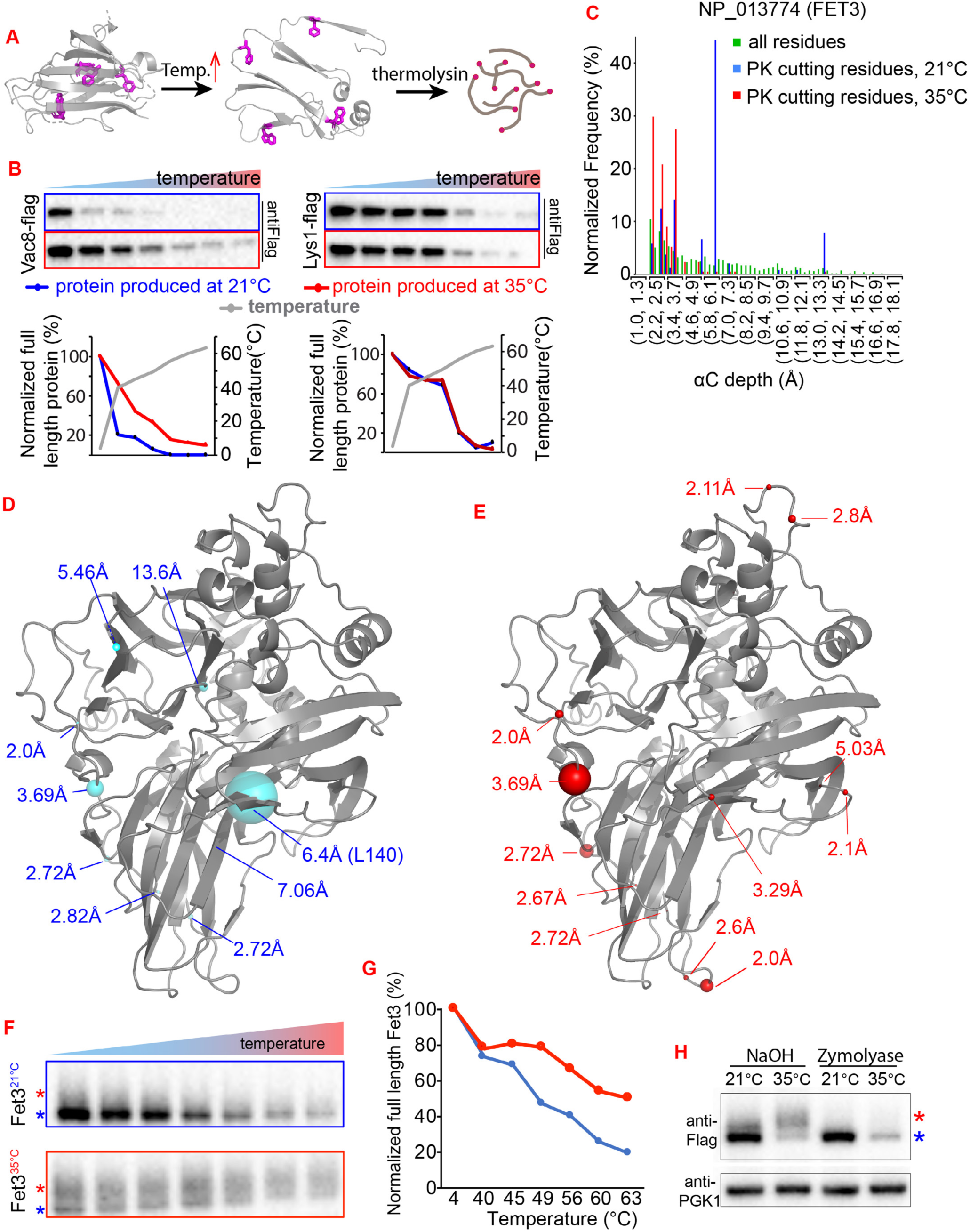
Temperature-dependent conformations and localizations of Fet3p. **A.** Schematic illustration of FastPP assay. Hydrophobic residues (purple) are buried inside the native protein. The increase in temperature relaxes the protein structure and exposes the hydrophobic residues that are selectively recognized and proteolyzed by thermolysin. The abundance of thermolysin-resistant full-length protein is proportional to the thermostability of this protein. **B.** Representative western blot and quantification of FastPP assay for two hits selected in 3D profiling analysis. FastPP was carried out on whole cell lysate for samples from both temperatures side by side for each protein. **C.** The distribution of PK sites for Fet3p from different temperatures. The distribution of alpha carbon depth of the entire structure is plotted in green as reference. **D, E.** The localization and depth of PK sites found in Fet3p from 21°C (blue) and 35°C (red) samples. The size of ball indicates the frequency of each PK site. See also the movie S1 for the map of PK sites in 3D structure. **F, G.** Representative western blot and quantification of Fet3p in FastPP assay. FastPP was carried out on whole cell lysate for samples from both temperatures. Blue and red stars indicate the major isoforms produced at 21°C and 35°C. Same for other figures. **H.** Representative western blot showing that the Fet3p produced at different temperatures have different migration patterns. Cell wall were digested with NaOH or Zymolyase. The protease in Zymolyase degraded the high molecule weight species. Images are representative of at least two independent experiments. See also Figure S5.

Finally, to investigate the possibility that conformational differences were caused by mutation, cells from both temperatures were subjected to whole-genome sequencing and then compared for differences between conditions. No mutations were identified within the coding regions of these 61 proteins. Among the 6 samples (3 from each temperature), we found 49 variant loci with an average depth of 42 reads, and none of these loci exhibit a uniform allelic difference between 21°C and 35°C (Table S5). In addition, ploidy analysis reveals no variation in chromosomal copy number between the strains (data not shown, see Method for details). Taken together, these results imply that, for these 61 proteins, the same polypeptides exist in stable alternative conformations at two temperatures.

### Temperature-dependent folding and thermostability of Fet3p

One of these candidates, Fet3p, was particularly interesting because its localization also differs between the two temperatures: at 35°C the protein localizes to the plasma membrane (PM), whereas at 21°C, ∼80% of the protein localizes to the ER with the rest in the PM (Singh et al., 2006; Stearman et al., 1996)(Figure 2A, 6A). The differences in protein structure, thermostability, and subcellular localization at two temperatures prompted us to focus on Fet3p to interrogate the functional consequence of these differences, if any.

**Figure 6:**
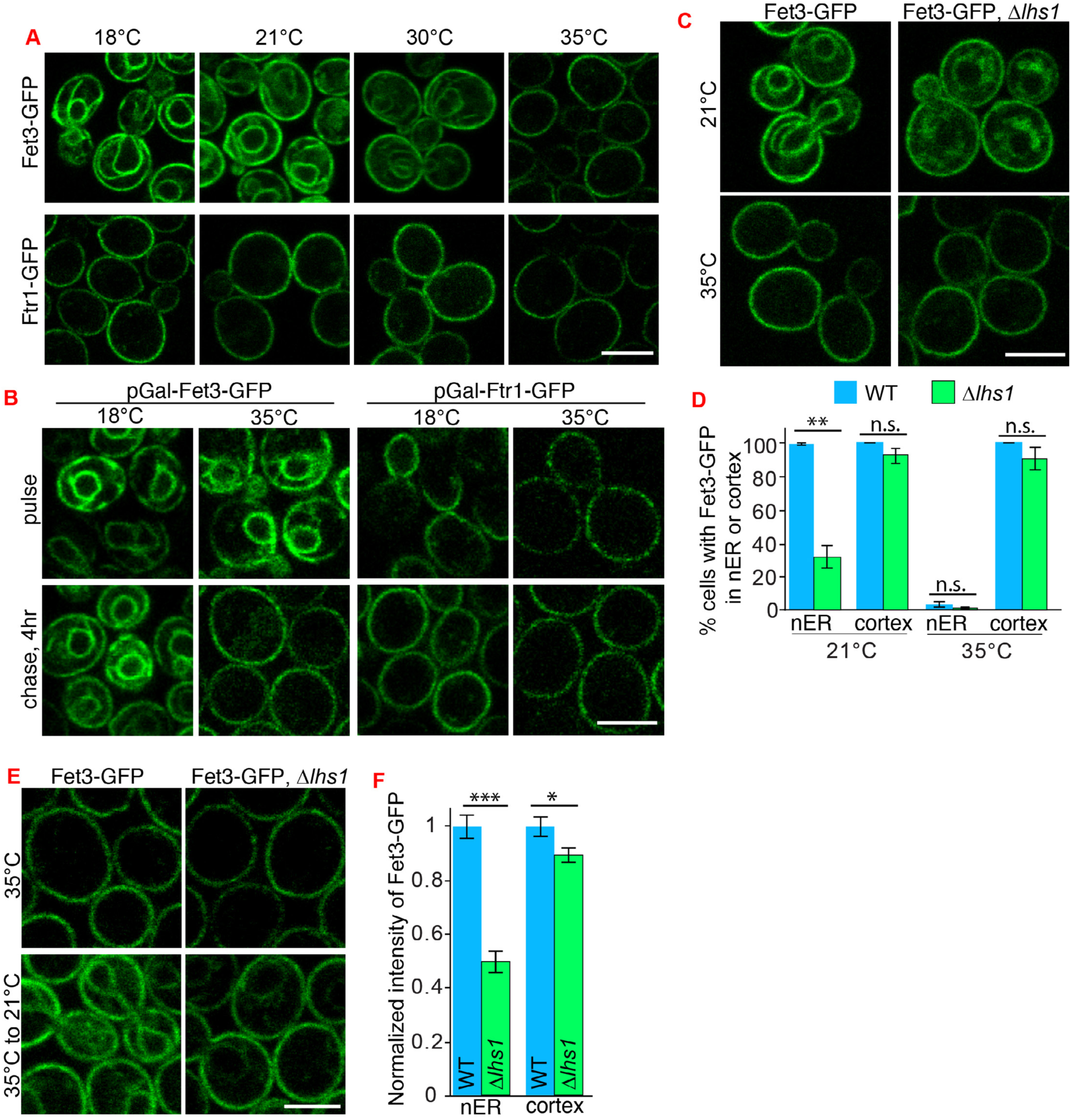
Fet3p localizes to ER at 21°C and requires the ER chaperone Lhs1p. **A.** Representative images of Fet3-GFP and Ftr1-GFP at different temperatures. Cells were cultured and refreshed at the indicated temperatures for 22 hours. **B.** Representative images of pulse-chase experiment for Fet3-GFP and Ftr1-GFP. Cells were cultured in YEP supplemented with Raffinose for more than 18hrs at indicated temperature before Gal induction. After 2hr Gal induction (pulse), glucose was added to terminate the Gal induction and the induced proteins were chased for another 4 hours. **C, D.** Representative images and quantifications of Fet3-GFP in wild type and *Δlhs1* cells that grown at indicated temperatures for 18 hours. Bar graphs are the normalized mean and SEM from three independent repeats. **E, F.** Representative images and quantifications of Fet3-GFP in temperature shift experiment for wild type and *Δlhs1* cells. Cells cultured overnight at 35°C were shifted to 21°C for 7hrs before images were taken. More than 50 cells were quantified for each strain. Scale bars are 5μm. Data were analyzed with unpaired two-tailed t test: ***, p<0.001; **, p<0.01; *, p<0.05; n.s., not significant. Images are representative of at least two independent experiments. See also Figure S6.

The PM localization of Fet3p at 35°C is consistent with the reported localization and function of Fet3p as a ferroxidase (Figure 2A, 6A) (De Silva et al., 1995; Stearman et al., 1996). The PK accessible sites at 35°C were mapped to the surface of the Fet3p reference structure, indicating that the available structure is a good reference for Fet3p at 35°C (Figure 5C, E, Movie S1). In contrast, based on the reference structure, the major PK sites at 21°C are predicted to be located inside the protein (Figure 5C, D, Movie S1). As indicated before, this is likely not due to differential conformational breath of the same structure at different growth temperatures, because the limited proteolysis was done by unifying the temperatures of samples to 4°C and later to room temperature (23°C). In addition, at 21°C, proteinase K could access some internal residues of the Fet3p reference structure more easily than the residues located on the surface of the reference structure (Figure 5D, E), suggesting a certain structural rearrangement of Fet3p made at 21°C that masks some surface residues and also stably exposes some residues that are buried in Fet3p made at 35°C.

The Fet3p produced at different temperatures showed different mobilities in denaturing gel (Figure 5H). Protease protection assay indicated that the higher molecular weight species are exposed to the extracellular space, representing the Fet3p on the PM (Figure 5H), since ER resident Fet3p is inaccessible to the external protease. The ratio of protease sensitive and resistant Fet3-flag is consistent with the observed subcellular distribution of Fet3-GFP between PM and ER (Figure 5H, 6A). When denatured and treated with Endo H, a highly specific endoglycosidase, Fet3p from different temperatures migrated as a single band of similar molecular weight, indicating that differential glycosylation is responsible for the mobility differences (Figure S5E). FastPP showed that Fet3p made at 35°C is more thermostable than Fet3p made at 21°C (Figure 5F, G, S5B). Thermolysin did not cleave the Fet3p at 4°C or at room temperature, but quickly digested the protein after thermal denaturation (Figure 5F and data not shown), indicating that there was no preexisting misfolded Fet3p before thermal denaturation. Thermostability difference persisted after de-glycosylating the purified Fet3-flag by Endo H, albeit to a lesser extent (Figure S5F), indicating that glycosylation unlikely accounts for differential thermostability. Although conformation changes do not necessarily cause thermostability changes, the thermostability differences of the purified and de-glycosylated Fet3p suggest that the proteins produced at different temperatures have different sets of noncovalent bonds. Therefore, consistent with the 3D profiling result (Figure 5C-E), the thermostability difference of Fet3p is rooted in the protein conformational difference and likely accentuated by glycosylation.

### ER-resident Fet3p (ER-Fet3p) is not an intermediate of plasma membrane Fet3p (PM-Fet3p)

By shifting temperature back and forth, we found that Fet3p distribution is tunable and dependent on growth temperature: the lower the environmental temperature, the more Fet3p localizes to the ER (Figure 6A). Fet3p is known to be a ferroxidase and associates with Ftr1p to facilitate the trans-membrane import of irons from the extracellular space (Singh et al., 2006; Stearman et al., 1996). Fet3p and Ftr1p rely on each other for export from the ER to the PM and knockout of one traps the other in the ER (Stearman et al., 1996). Thus, we asked whether the localization of Fet3p in the ER at low temperature is due to the lack of Ftr1p or export defect of Ftr1p. However, we found Ftr1p localizes to PM at all tested temperatures (Figure 6A, S6A).

As Fet3p expressed at two different temperatures has distinct glycosylation states, we asked whether the ER localization of Fet3p was due to general glycosylation defects at low temperature. We examined the localization of multiple plasma membrane proteins containing similar N-glycosylation and O-glycosylation sites as Fet3p (Neubert et al., 2016; Zielinska et al., 2012) and found that none of these plasma membrane proteins localized to ER at low temperature (Figure S6B). To test whether the ER-Fet3p at low temperature is an intermediate that will be processed and shipped to the plasma membrane over time, we performed a pulse-chase assay to follow the Fet3-GFP expressed at both temperatures. The kinetics of GAL-induced expression of Fet3-GFP were similar at both temperatures (Figure 6B). While Ftr1-GFP at both temperatures was able to migrate to plasma membrane, the Fet3-GFP expressed at 18 °C, but not at 35 °C, stayed in ER even after extended chase (Figure 6B). Moreover, newly synthesized Ftr1-GFP was asymmetrically deposited in daughter cells, while Fet3-GFP was deposited in both mother and daughter cells at low temperature, indicating that these two proteins were processed differentially at low temperature (Figure S6C). Therefore, the ER localization of Fet3p at low temperature was not due to defective Ftr1p, glycosylation, or export from the ER.

### ER chaperone Lhs1p is required to produce ER-Fet3p

After ruling out possible ER defects, next we asked how does the same Fet3p protein achieve distinct temperature-dependent localizations. Given the conformational difference between Fet3p produced at different temperatures, we wondered whether ER chaperones may play a role in regulating the differential localization of Fet3p. Deletion of most ER-resident chaperones had no effect on Fet3p localization (Figure S6D), except the deletion of LHS1. More than 65% *Δlhs1* cells lost ER-Fet3p at low temperature, while Fet3p localization to the plasma membrane was largely unaffected (Figure 6C, D). This is not a general problem that affects all ER proteins as ER protein Sec63-GFP showed a similar distribution in both wild type and *Δlhs1* cells (Figure S6E). Moreover, when the cells were switched from high temperature to low temperature to induce the ER-Fet3p in wild type cells, *Δlhs1* cells failed to generate as much ER-Fet3p as in wild type cells (Figure 6E, F). As the functions of chaperones are to eliminate misfolded proteins, the lack of ER-Fet3p, instead of accumulating more Fet3p in ER, in *Δlhs1* cells suggests that ER-Fet3p are not misfolded proteins and that at lower temperatures, cells actively produce a pool of ER-localized Fet3p with the help from chaperone Lhs1p.

### Identification of ER proteins that interact with ER-Fet3p

To understand the function of Fet3p in the ER, we purified the Fet3p from cells grown at 21°C and 35°C and used mass spectrometry to characterize temperature-specific protein-protein interactions (PPIs) (Table S6). We found five unique ER-resident proteins that co-purified with Fet3p at low temperatures and three of these proteins (Scs2p, Skn1p, Elo3p) are related to lipid metabolism. Similar enrichment of lipid metabolic proteins was also observed in the list of Fet3p-interacting proteins from previous large-scale screens, which were characterized at temperatures with some Fet3p localizing to the ER (Figure 7A, Table S6) (Miller et al., 2005). Co-IP confirmed that the interaction between Fet3p and Elo3p, the common hit between our and previous large-scale analysis, is temperature-dependent (Figure S7A).

**Figure 7:**
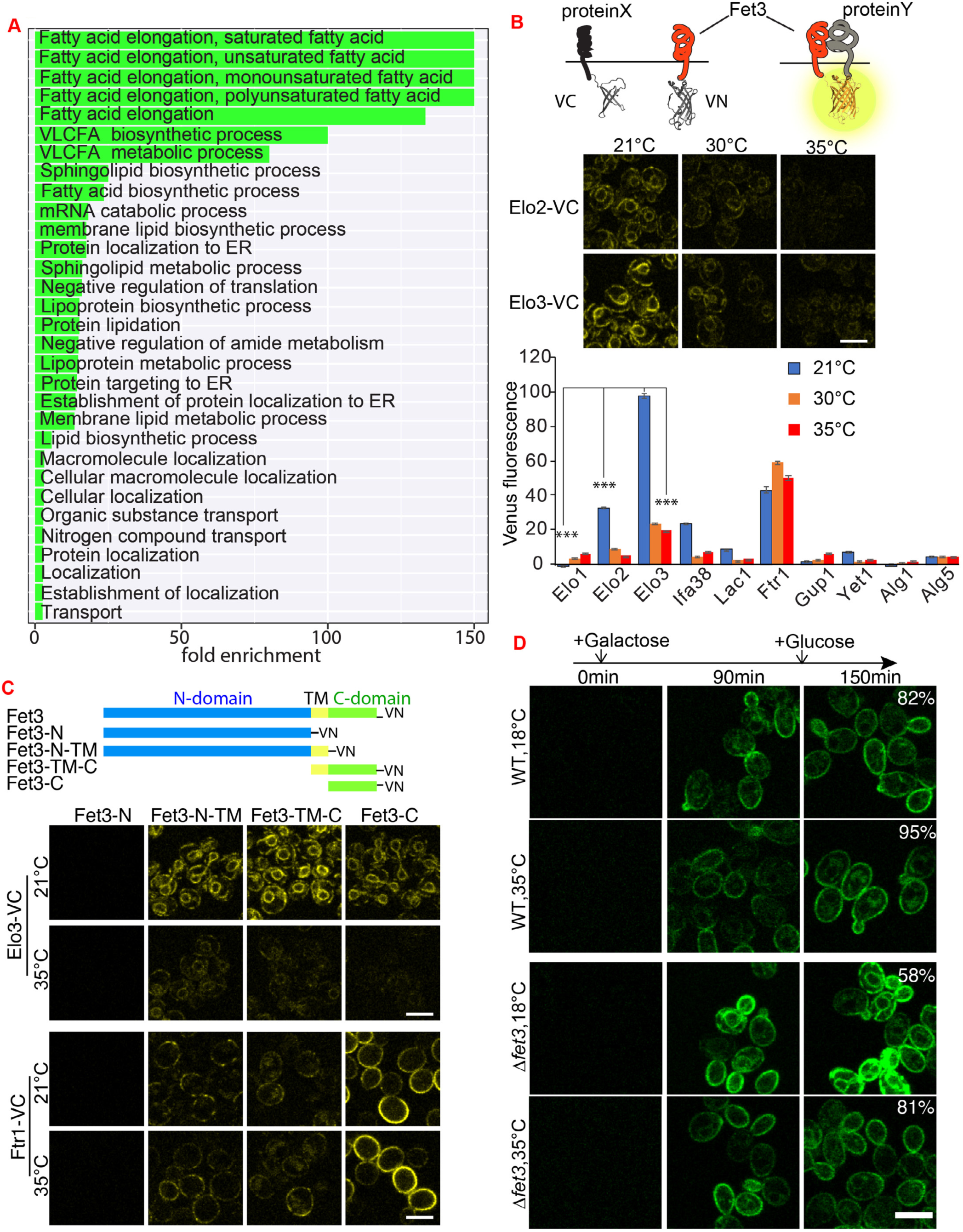
The cellular function of ER-localized Fet3p. **A.** GO term analysis of the proteins physically interact with Fet3p (Ashburner et al., 2000; Gene Ontology Consortium, 2021). **B.** Schematic illustration, representative images, and quantification of split Venus PCA assay. Interaction between Fet3p and candidates reconstitutes Venus fragments (VN and VC: N- and C- fragment of Venus). Signals from the cells with only Fet3-VN were used as control and were subtracted for each candidate at corresponding temperature. Shown is mean and SEM quantified from more than 100 cells. Data were analyzed with unpaired two-tailed t-test: ***, p<0.001. **C.** Schematic illustration and representative images of interactions between different Fet3p fragments and full length Elo3p or Ftr1p revealed by Venus PCA assay. N-domain, N terminal domain facing ER lumen; TM, transmembrane domain; C-domain, Cytosolic domain. VN and VC as in (B). **D.** Deleting FET3 retarded the maturation of Gas1. Gas1-GFP was pulse expressed from Gal promoter for 120min and traced for another 30min after adding glucose. Percentage of cells with PM-localized Gas1-GFP was quantified and shown for each strain. Only *Δfet3* cells grown at 18°C show ER and internal signals compared with the PM-localized Gas1-GFP in wild type cells and 35°C. Scale bars are 5μm. Images are representative of at least two independent experiments. See also Figure S7.

To complement the biochemical assays, we used a protein-fragment complementation assay (PCA) based on split Venus (Jin et al., 2011; Weill et al., 2018) (Figure 7B), where physical interaction between two proteins creates a fluorescence signal. Using this PCA assay, we found that Fet3p and Ftr1p interact both at 21°C and 35°C, but ER-resident lipid metabolic proteins, such as Elo3p, show strong fluorescence reconstitution specifically at 21°C (Figure 7B). These reconstitutions cannot be attributed to a difference in protein level of these ER proteins (Figure S7B).

We then asked which part of Fet3p is involved in the temperature-dependent interactions. Fet3p is a type-I transmembrane protein with N-terminal (N) domain, transmembrane (TM) domain, and cytosolic (C) domain (Figure 7C). The cytosolic domain of Fet3p interacts with Ftr1p, as reported previously (Singh et al., 2006) (Figure 7C). We found that the cytosolic domain is also required for Fet3-Elo2/3 interaction (Figure 7C, S7C). In addition, the TM domain of Fet3p seems to enhance the temperature-specific interaction between Fet3-Elo2/3, while it reduces the Fet3-Ftr1 interaction at both temperatures (Figure 7C, S7C). This indicates that the lower temperature and probably the changes of ER lipid composition (Figure S1F), allow the TM domain of Fet3p to adopt a conformation with higher affinity for Elo2/3p than Ftr1p. Interestingly, the Fet3-C fragment, which localizes to cytosol (Figure S7D), also interacts in a temperature-dependent manner with Elo2/3p (Figure 7C, S7C). Therefore, the interaction between Fet3p and ER proteins does not depend on ER-localization and the ER-associated glycosylation of Fet3p. As there is no post-translational modification difference for Fet3-TM and Fet3-C between these two temperatures (Table S4), these results suggest that the conformations of Fet3-TM and Fet3-C domains were shaped by the culture temperatures to regulate its PPIs and carry out temperature-specific functions.

### The function of ER-Fet3p

What are the functional consequences of the interaction between ER-Fet3p and the sphingolipid synthesis enzymes Elo2/3p? To address this question, we measured cellular sphingolipid composition in wild type and *Δfet3* cells. Elo3p is known for its role in extending the fatty acid chain from C24 to C26, and knockout of ELO3 has been shown to reduce the C26 level in several sphingolipids (Montefusco et al., 2014). We purified and analyzed the lipids from *Δfet3* and wild type cells cultured at different temperatures by LCMS/MS. This assay detected three sphingolipids that showed a significant reduction in *Δfet3* cells grown at 18°C, but not in cells cultured at 35°C (Figure S7E).

C26 fatty acids and sphingolipids are critical for the biogenesis of GPI-anchor proteins. The maturation of GPI-anchor proteins, such as Gas1p, is associated with the exchange of an acyl chain at the sn-2 with a C26 saturated acyl chain (Bosson et al., 2006). In addition, the sphingolipids are critical components of lipid rafts that are required for the efficient export of GPI-anchored proteins from ER (Mayor and Riezman, 2004). Therefore, we tested whether deleting FET3 affects the maturation of Gas1p. When Gas1-GFP was expressed transiently in wild type cells grown at 18°C, most of the newly synthesized Gas1p matured and migrated out of the ER (Figure 7D). However, the maturation of Gas1-GFP was delayed in some *Δfet3* cells at 18°C (Figure 7D). In contrast, Gas1-GFP matured and localized normally in *Δfet3* cells at 35°C. Taken together, these results suggest that, at low temperature, Fet3p plays a role in sphingolipid synthesis and ER homeostasis. As there are some additional PPIs of Fet3p not investigated (Figure 7A), it is possible that Fet3p is also involved in other functions in the ER.

## Discussion

To adapt, organisms need to resolve two issues: protect the processes that are essential for survival under any condition and generate new activities that are either necessary or better suited for the new environment. Sudden environmental changes cause proteins to misfold and aggregate, which are removed by protein quality control machinery. However, it is unknown whether these proteins get stuck in a futile cycle of synthesis-misfolding-degradation when the environmental change persists. Using proteome-wide analysis of yeast grown at two different temperatures, we find that cells employ different strategies in response to long-term environmental change: reducing the expression of some thermolabile proteins and alternative protein folding. The alternation of protein conformation is a new feature of thermo-adaptation. Similar protein conformation changes based on surface accessibility difference have been suggested for cells grown at different nutrient conditions or acute environmental stresses (Cappelletti et al., 2021; Feng et al., 2014). However, surface accessibility of a protein is defined by intrinsic conformation or inter-molecular interactions, which are difficult to distinguish (Cappelletti et al., 2021; Feng et al., 2014). Using reference structure-based analysis, we were able to partially dissect the causes of surface accessibility changes and found that for some proteins, changes in protein surface are linked to alternative protein conformation. In addition, we found that the temperature-specific conformations, localization, and protein complexes are associated with temperature-specific cellular functions. We postulate that the conformation-function plasticity exemplified by Fet3p to be a general theme for other alternative folding proteins identified in this work. What remains to be determined is the extent of the conformational differences and the exact nature of the conformational differences. In the future, detailed structural studies of individual proteins isolated from the two temperatures should address these issues.

There are sporadic examples of the same protein adopting different structures and functions, so-called metamorphic proteins (Murzin, 2008). The studies of metamorphic proteins indicated that protein folding states can be context dependent. For example, different native folded states of the Lymphotactin are shaped by the environment to perform distinct functions, while delta-Crystallin exists as a structural protein in the eye lens and as an active arginosuccinate lyase in other cell types (Tuinstra et al., 2008; Piatigorsky et al., 1988). Our findings suggest that the environment-dependent conformation change is likely part of a much broader phenomenon (Figure S7F). Conformational plasticity, which allows functional changes without changing primary sequence, can expand the functional proteome space and could serve as an adaptative tool in changing environments.

What cellular processes restrict or allow proteins to switch conformation? More specifically, how does Fet3p acquire different conformations, and thereby functions, at different growth temperatures? Some hints can be found from other alternative folding and metamorphic proteins. For example, the synonymous mutations on MDR1 genes change the decoding speed of some residues and the folding path of the polypeptide, which folds as a drug-resistant conformation in some cancer cells (Kimchi-Sarfaty et al., 2007). Although none of the 61 proteins with alternative conformations identified in our study contains single nucleotide polymorphisms (SNPs), the thermal acclimatization modifies tRNA biogenesis and modifications (Figure 1E, F), which in turn can change the decoding speed and folding paths of some proteins (Pechmann and Frydman, 2013; Pechmann et al., 2014). In another example, cytosolic and mitochondrial aconitase acquire different structures and functions through the binding of a co-factor: while the association of cubane iron sulfur cluster (ISC) with aconitase is required for its enzymatic activity towards tricarboxylic acid (TCA) cycle substrates, the loss of this cubane cluster causes conformational changes in aconitase and transforms it into a mRNA binding protein (Klausner and Rouault, 1993). Similar to aconitase, Fet3p that migrates through the secretory pathway together with Ftr1p to the PM is known to contain 4 Cu ions as co-factors gained at the post-Golgi apparatus (Taylor et al., 2005). We hypothesize that the folding of Fet3p into a ferroxidase that is recognizable by Ftr1p and sorting factors is sensitive to the environmental temperature and the ER environment. At higher temperatures, the Fet3p attains a structure that is recognized by Ftr1p and sorting factors, and thus migrates through the secretory pathway, gains access to the Cu ions in post-Golgi apparatus and matures into an active ferroxidase. At lower temperatures, most of the Fet3p polypeptides acquire a different folding state that cannot be recognized by Ftr1p and sorting factors. The precise mechanism regulating the production of ER localized Fet3p at low temperature requires further study. Our preliminary findings indicate that a specific chaperone, Lhs1p, is important to change Fet3p’s folding state. In fact, the role of protein chaperones in generating phenotypic variation has been observed previously. For example, Hsp90p, when attenuated, releases otherwise silent protein variants and generates new traits (Jarosz and Lindquist, 2010; Rutherford and Lindquist, 1998). We find that thermal adaptation requires the activity of specific chaperones. This is a departure from the view that chaperones restrict conformational variation to chaperones promoting conformational variation. Further studies should provide insight into this previously unanticipated possibility.

## Acknowledgements

The authors thank E. Verdin, G. Lithgow, B. Kennedy, P. Walter, P. Haghighi, and R. Brem for discussion, T. Payne and A. Holtz for assistance in mass spectrometry sample preparation, A. Lawlor and M. Peterson from Sequencing and Discovery Genomics Team for NGS. This work is supported by T32-AG052374 to M.Domnauer, SIMR to K.Si, and DP5OD024598 to C.Zhou, the NCRR shared instrumentation grant 1S10 OD016281 to the Buck Institute.

## Author contributions

Conceptualization, C.Z. and K.S.; investigation, C.Z., D.M., F.Z., L.L., Y.Z., C.E.C., J.R.U., J.C., S.M., L.F., Y.Z., C.S., B.S., and R.S.; resources, C.Z., K.S., A.R.; software, C.Z., L.L., Y.Z., J.R.U., B.F.; supervision, C.Z. and K.S.; writing, original draft, C.Z. and K.S.; writing– review and editing, all authors.

---

**Figure S1:**
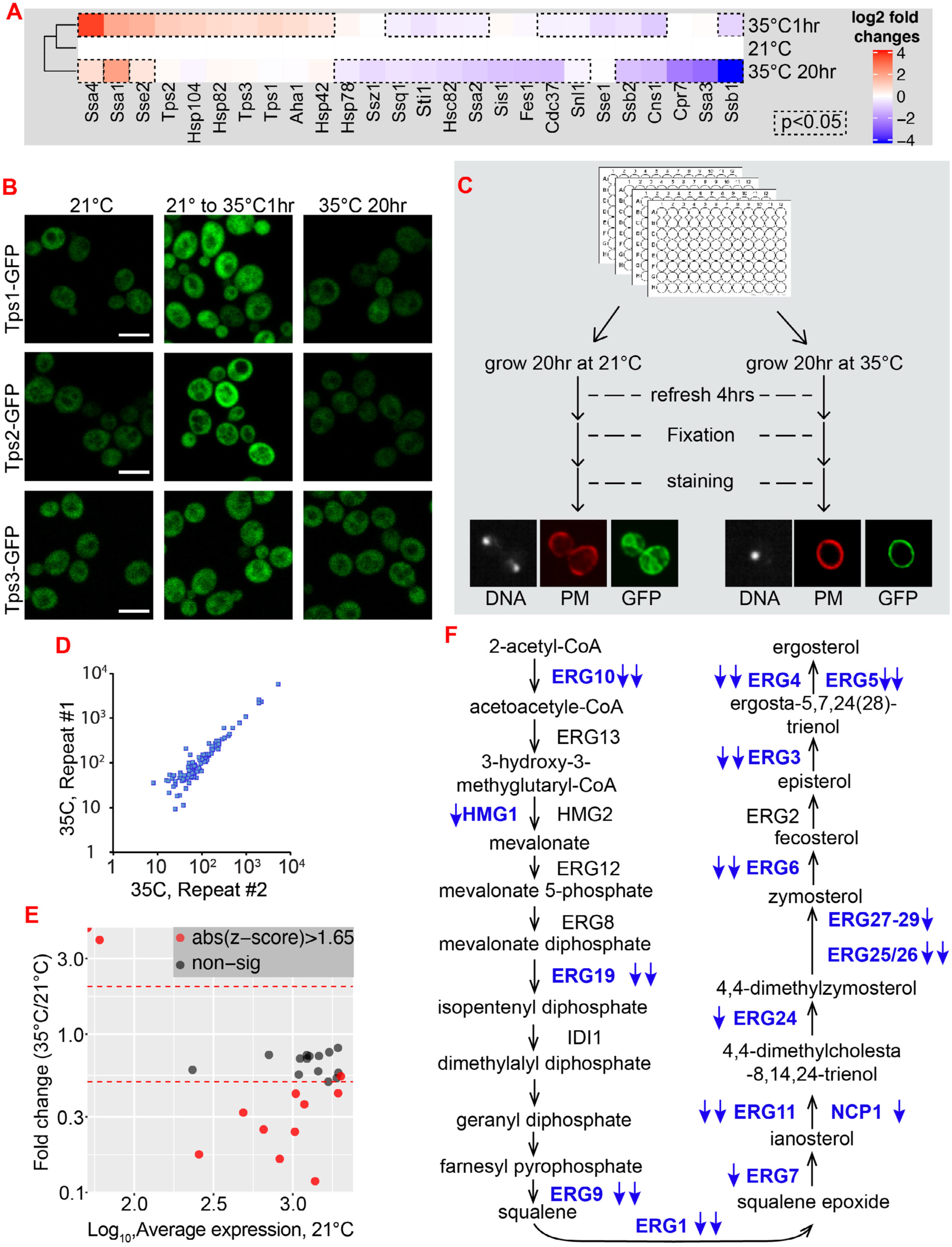
Expression changes of different proteins in response to acute and chronic temperature stress, related to Figure 1. **A.** Expression level of different chaperones in cells that experienced transient heat shock (1 hour shift from 21°C to 35°C) or long-term high temperature acclimation. The expression levels were normalized to cells grown at 21°C for each chaperone. Shown is mean and SEM from more than 50 cells for each sample. **B.** Representative fluorescence images of cells expressing GFP tagged Tps1/2/3p grown at different conditions. Scale bar: 5μm. **C.** Design of GFP-based screening to assess proteome-wide changes during thermal acclimation. DNA and plasma membrane (PM) were stained by Hoechst dye and FM4-64, respectively. **D.** Quantification of protein levels from two replicates of a testing plate during the GFP screen. This suggests that the screening protocol is stable and reproducible. **E.** Expression level of proteasome proteins at different temperatures from the GFP screen. Red lines indicate the 2 folds increase or decrease. **F.** The expression changes of key enzymes in ergosterol synthesis pathway. Blue fonts indicate reduced expression and double arrows indicate more than 2 folds reduction. Images in this figure are representative of at least two independent experiments.

**Figure S2:**
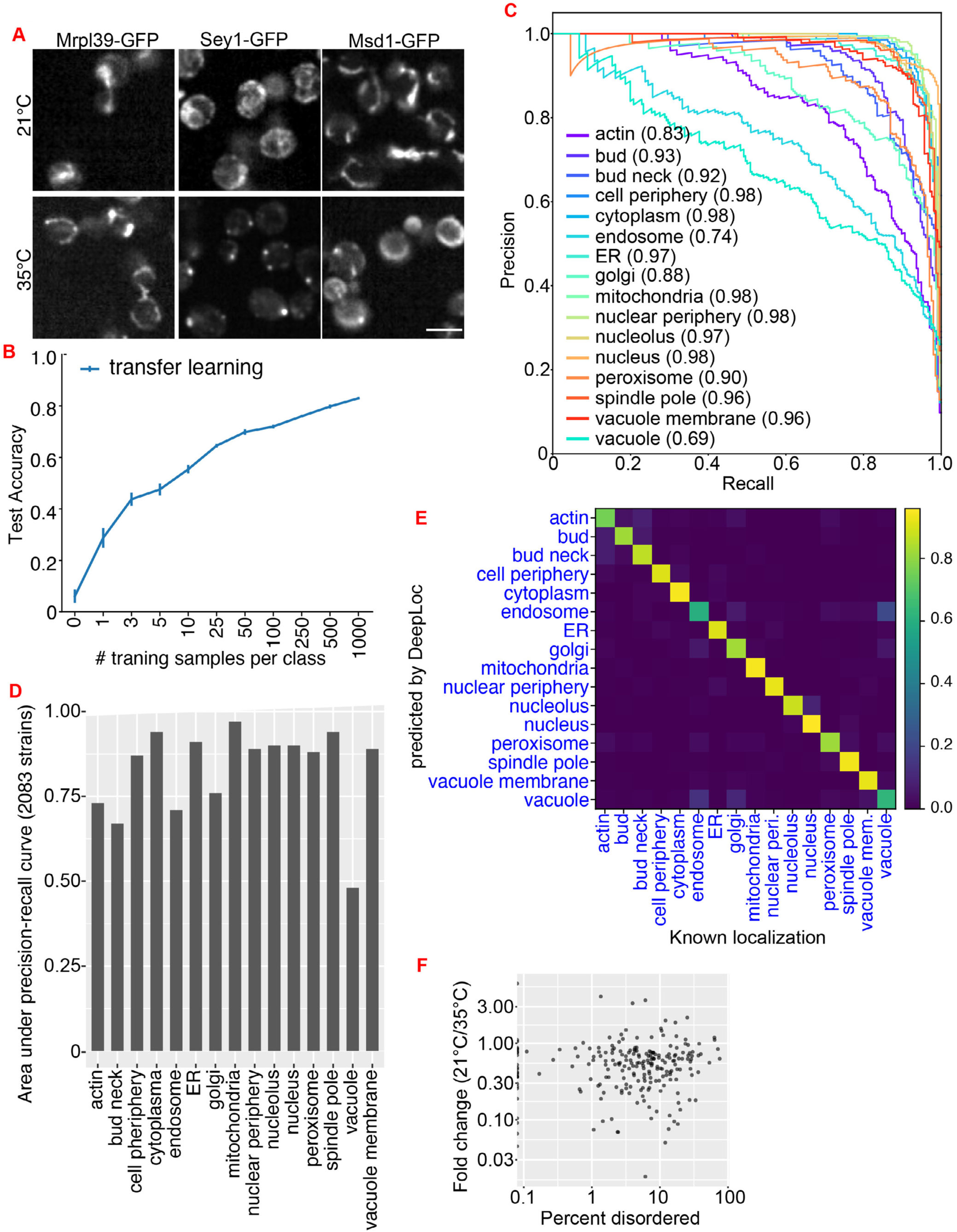
Additional data from the training and evaluation of the DeepLoc model, related to Figure 2. **A.** Example of proteins showing cellular localization changes during thermal adaptation. Scale bars: 5μm. **B.** Relationship between the number of training images per class vs classification accuracy during transfer learning of DeepLoc. For each size of training datasets, we run 5 batches of randomly sampled images. The error bars are the standard deviation of the classification accuracy from the 5 runs. **C.** Precision-Recall curve for each localization class, based on the prediction of 3866 single cell (test data set). Numbers in the bracket indicate the Area Under Curve (AUC) of each localization class. **D.** Calculated Area Under Curve (AUC) for each localization class, based on the prediction of strains annotated to have a single localization in (Huh et al., 2003). **E.** Confusion matrix based on 3866 single cell tests for the model after transfer training with 1,000 training images per class. For each cell, the localization class that gets the highest prediction score was set as the predicted localization (y-axis). X axis is the annotated subcellular localization for each cell. **F.** No correlation between the expression change and intrinsic disorder of these aggregation prone proteins.

**Figure S3:**
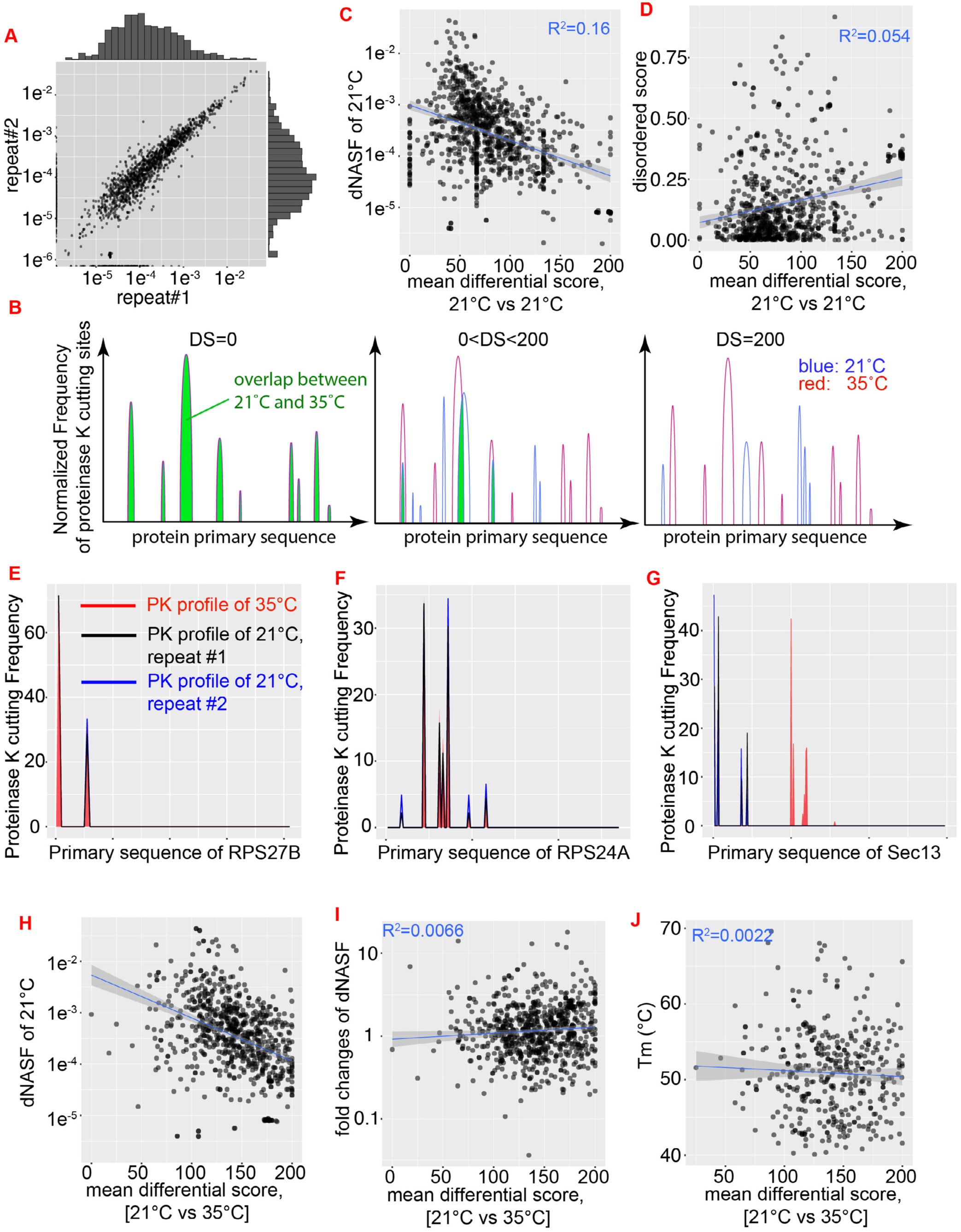
Additional analysis for the data from limited proteolysis mass spectrometry, related to Figure 3. **A.** The mass spectrometry results were highly reproducible between different repeats. The distributed Normalized Spectral Abundance Factors (dNSAF) of two (out of three) repeats from 21°C samples were plotted against each other. **B.** Schematic illustration of differential score (DS) defined by the difference between PK profiles of different samples, either from biological repeats or different temperatures. The DS is calculated by first normalizing the frequency of PK cutting for each residue to the total PK cutting events for a given protein, therefore, the total underlying area for each red and blue profile is 100. This is followed by subtracting the normalized frequency of each residue between two samples of the same protein and aggregating the absolute value of difference from each residue to give rise to the final differential score. Zero differential score between two 1D-profiles means proteinase K cuts the same surface residues with the same frequency, thus, indicates that this protein has the same surfaces in different samples; a differential score of 200 means the cutting sites are completely different in different samples, which suggests completely different surfaces exist for this protein at different conditions. **C.** Limited correlation between protein abundance (dNASF) and the noise of 1D profiles between biological repeats of 21°C. **D.** Limited correlation between the predicted disordered scores of each protein and its noise in 1D profiles of 21°C biological repeats. Disordered scores were predicted by DisoPred3 program (Jones et al., 2015). **E, F**. Example of proteins with no significant 1D profile shift between temperatures. **G**. Example of protein with a significant 1D profile shift during temperature adaptation. **H-J**. Limited correlation between protein abundance (H), expression changes (I), melting temperature (J) and 1D profile shifts as measured by differential scores between two temperatures. Melting temperatures were from previous publication (Leuenberger et al., 2017).

**Figure S4:**
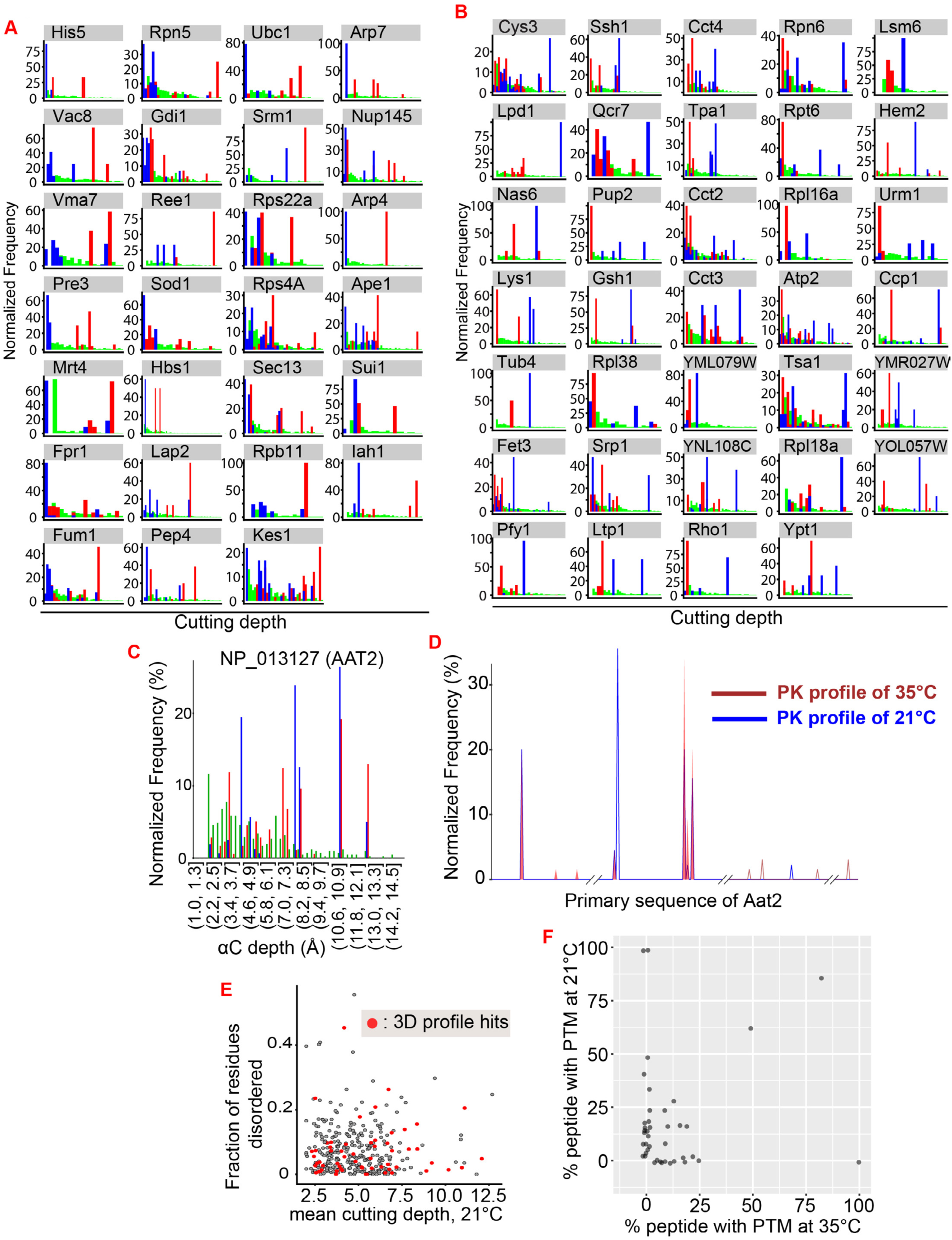
3D profiles of 61 hits, related to Figure 4. **A.** Additional examples for Figure 4E showing that the PK sites observed in the 35°C samples are mapped deeper than in the 21°C samples in reference structures. The distribution of alpha carbon depth of the entire structure is plotted in green as reference. **B.** Additional examples for Figure 4F showing that the PK sites in the 21°C samples are mapped deeper than in the 35°C samples in reference structures. The distribution of alpha carbon depth of the entire structure is plotted in green as reference. **C.** Example of a protein with PK sites mapped deeply into the reference structure at both temperatures. The distribution of alpha carbon depth of the entire structure is plotted in green as reference. Note that the distribution of PK cutting depth is different from the overall distribution of all residues in this protein. The enrichment of the cutting sites deeply into the reference structure indicates that the proteinase K has access to some inner residues more easily than the residues located on the surface of the reference structure. See also (**D**) and movie S1 for the map of cutting sites in 3D structure. **D.** 1D profile for the protein shown in (**C**). Note that the cutting sites are strongly enriched to certain regions and not widespread throughout the entire protein, indicating that the protein was not unfolded and randomly digested by proteinase K. **E.** Limited correlation between the mean cutting depth of 21°C samples and the predicted disordered score for each protein. The proteins with temperature-driven cutting depth changes (cyan dots in Figure 4I) are colored in red. **F.** Percentage of each peptide that contains PTM. We observed 44 peptides with PTM for 20 out of 61 proteins. Total distributed spectrum of each peptide with and without PTM were pooled from three replicates of each temperature to calculate the percentage.

**Figure S5:**
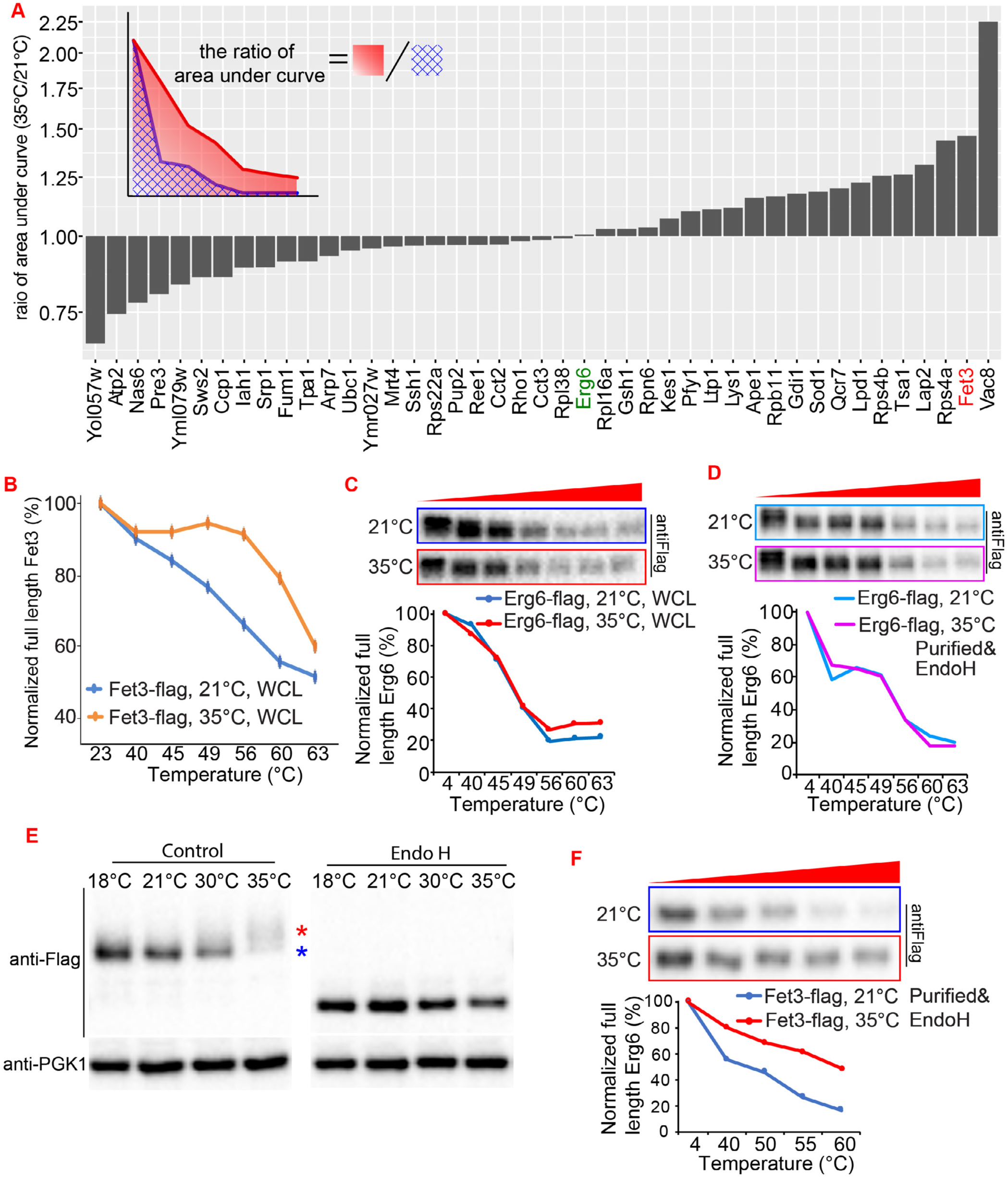
FastPP analysis of different hits, related to Figure 5. **A.** The thermal stability difference of proteins produced at 21°C and 35°C represented by the difference of area under curves of the FastPP assays between different temperatures. Hits from Figure S4A and B were C-terminally tagged with 3xflag and cultured at different temperatures. Some of the hits resisted flag tagging and were not included. Fresh cells from both temperatures were lysed and the FastPP was carried out on whole cell lysate (WCL) for samples from both temperatures side by side for each protein. At least two repeats of FastPP assay were done for each protein and the average ratio of area under curve from two temperatures were shown. **B.** FastPP curve of Fet3-flag in whole cell lysate lysed with 0.5% triton X100 instead of 0.34 mM n-Dodecyl-B-D-Maltoside (DDM), which was the detergent used for all other FastPP assays. **C, D.** Western blot and quantification of FastPP assay for Erg6-flag in whole cell lysate (**C**) or purified/Endo H treated (**D**). **E.** Western blot of the Fet3-flag produced at different temperatures. The same number of cells from different temperatures were lysed and treated with EndoH to remove glycosylation. **F.** Western blot and quantification of FastPP assay for Fet3-flag that was purified and treated with Endo H to remove glycosylation difference. For all FastPP assays, at least two repeats were done, and the representative results are shown.

**Figure S6:**
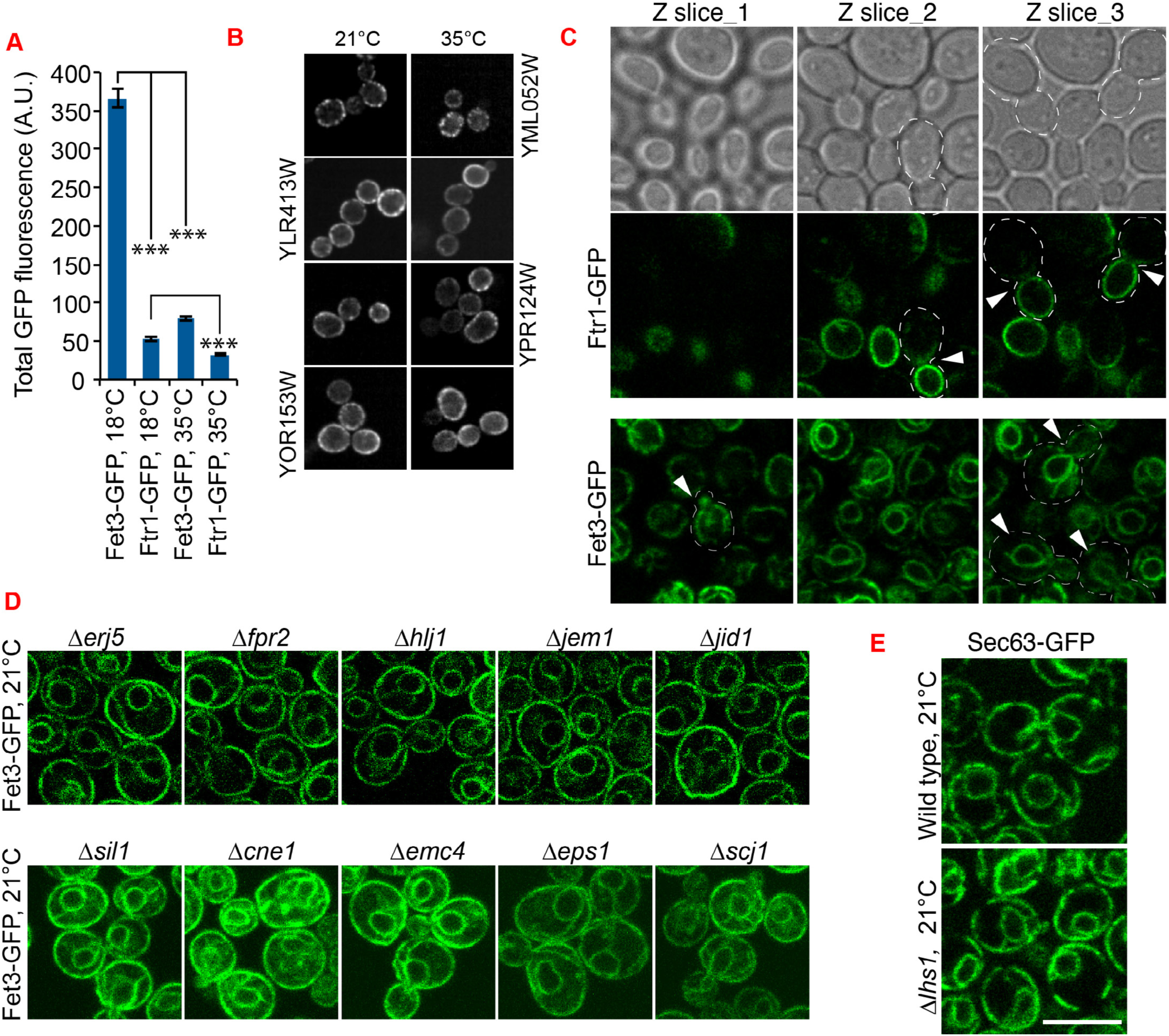
Additional data for the role of LHS1 in regulating ER-Fet3, related to Figure 6. **A.** Expression level of Fet3-GFP and Ftr1-GFP at different temperatures. The z-stacks of cells grown at different temperatures were sum projected and the total amount of GFP fluorescent signal was measured for each cell. Shown is mean and SEM from more than 100 cells for each condition. Data were analyzed by unpaired two-tailed t-test: ***, p<0.001. **B.** Cellular localization of other glycosylated plasma membrane proteins at different temperatures. Images from GFP screen. **C.** Representative subcellular localization of Ftr1-GFP and Fet3-GFP in cells grown at 18°C during pulse chase experiment. White arrow heads point to the cells showing featured protein distribution across bud-mother axis. Note that Ftr1-GFP localized to daughter cell and the Fet3-GFP localized to both mother and daughter cells. All scale bars in this figure are 5μm. **D.** Cellular localization of Fet3-GFP in different chaperone mutants. **E.** Cellular localization of Sec63-GFP in wild type and *Δlhs1* mutant cells. Images in this figure are representative of at least two independent experiments.

**Figure S7:**
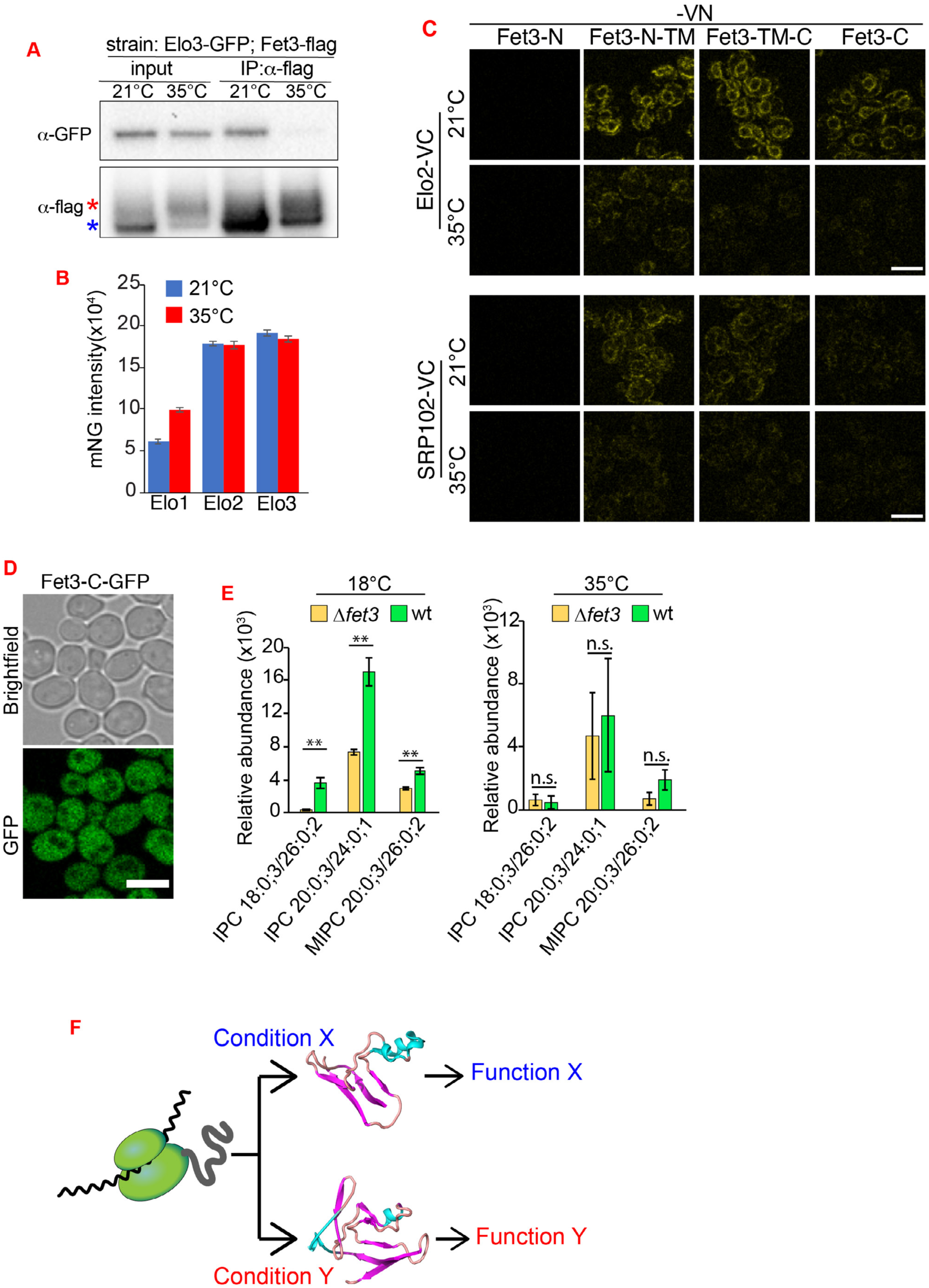
Temperature-specific functions of Fet3p, related to Figure 7. **A.** Co-IP of Elo3-GFP with Fet3-flag at different temperatures. Same amount of cells were used for both temperatures. **B.** Expression level of Elo1/2/3p quantified from images. Elo1/2/3 were tagged with mNeonGreen (mNG) from the C-SWAT library (Weill et al., 2018; Meurer et al., 2018). **C.** Venus PCA assay for different fragments of Fet3-VN with Elo2-VC or SRP102-VC. **D.** Subcellular localization of Fet3-C fragment visualized by GFP fused to its C terminus. **E.** Sphingolipid profiles of wild type and *Δfet3* cells from different temperatures. Shown are mean and SEM from three repeats. Data were analyzed by unpaired two-tailed t-test: **, p<0.001; n.s., not significant. **F.** Model of protein alternative folding under different environmental conditions. The nascent polypeptide can fold into different structures depending on the environment, which can directly affect the free energy of the folding path or through modifying the translation speed and PPIs of the proteins. All scale bars in this figure are 5μm. Shown is mean and SEM from three repeats. Images in this figure are representative of at least two independent experiments.

## STAR Methods

### RESOURCE AVAILABILITY

#### Lead Contact

Further information and requests for reagents and resources should be directed to the Lead Contact Chuankai Zhou.

#### Material Availability

All plasmids and strains generated in this work can be requested from the lead contact’s lab. All antibodies and chemicals used in this study are commercially available.

#### Data and Code Availability

NGS raw data has been deposited (https://www.ncbi.nlm.nih.gov/sra/PRJNA699272). The mass spectrometry datasets have been deposited to the MassIVE and Proteome Xchange repositories with the following accessions (password: winter): MSV000086790/PXD023924 for the MudPIT LiPMS analyses, MSV000086791/PXD023925 for the MudPIT PTMs analyses, and MSV000086243/PXD021862 for the Fet3-FLAG IP analysis. Original data underlying this manuscript can be accessed from the Buck Institute link at https://pubdata.buckinstitute.org/771146.html (Username: reviewer Password: 2yb81j6x) and the Stowers Original Data Repository at www.stowers.org/research/publications/LIBPB-1597 All codes are available upon request.

#### Experimental Model and Subject Details

All *S. cerevisiae* strains and plasmids used are listed in the Key Resources Table. Yeast strains used in this study are based on the BY4741 strain background. Genetic modifications were performed with PCR mediated homologous recombination (Longtine et al., 1998) and genotyped with PCR to confirm correct modification and lack of aneuploid for the chromosome that gene of interest is located. The GFP-tagged strains were from the GFP collection (Huh et al., 2003). The flag-tagged strains were generated by PCR amplification of 3xFlag cassette from tagging plasmids based on pFA6a (Funakoshi and Hochstrasser, 2009). The mutant strains were generated by PCR amplification of KanMX cassette from existing mutant strains (Yeast Knockout Collection, GE Dharmacon) or pFA6a-KanMX plasmid to transform By4741 or the strains based on By4741. Expression of proteins from integration plasmids was done by integrating the linearized plasmid into TRP1 locus. The Venus PCA strains were constructed using C-SWAT library and plasmids kindly provided by Dr.Knop and Dr. Schuldiner (Meurer et al., 2018; Weill et al., 2018). Anti-flag and anti-GFP are from sigma, A2220 and SAB4301138, respectively.

Yeast cells were grown in YEP with 2% dextrose (YEPD) for over 18hrs at indicated temperature. Pre-warmed or chilled fresh YEPD was used to refresh the culture for additional 4hrs before imaging. The OD_600_ of cell culture was maintained between 0.3-1.0 throughout the experiments and additional refresh was done when longer incubation is indicated. For some strains with low GFP expression, one wash with SC complete media was done before imaging to reduce the autofluorescence background from YEPD. To express proteins under Gal promoter, cells were grown in YEP or SC complete media containing 2% raffinose to OD_600_∼0.8 and induced by add 2% galactose for 2hr. Gal induction was stopped by adding 2% glucose for indicated time before imaging. All images were processed by Image J software (NIH, Bethesda, MD). For visualization purposes, images scaled with the bilinear interpolation were used for figures. For Ssa1-GFP, cells were cultured in YPED at indicated temperature for 18hrs and refreshed for another 6hrs. For Tps1-GFP, Tps2-GFP, and Tps3-GFP, the overnight culture was refreshed by adding glucose to 2%, followed by dilution with fresh, pre-warmed or chilled medium to desired OD. Imaging plates (Eppendorf # 0030741030), instead of regular glass slides and coverslips, were used for imaging. All medium was prepared by autoclave the solution without glucose/raffinose/galactose for 20min and add filtered carbon source as indicated.

### METHOD DETAILS

#### Confocal microscopy

Live images were acquired using a Carl Zeiss LSM-510 Confocor 3 system with 100x 1.45 NA Plan-Apochromat objective and a pinhole of one airy unit. The system was driven by Carl Zeiss AIM software for the LSM 510. 488/561 nm excitation was used to excite GFP/RFPs, and emission was collected through the appropriate filters onto the single photon avalanche photodiodes on the Confocor 3. All GFP images were acquired through a 500-550 nm filter. RFP images were acquired with a 580nm long pass filter on the Confocor 3. All images were acquired in a multi-track, alternating excitation configuration so as to avoid GFP bleed-through.

#### GFP screen for thermal adaptation

GFP strains were transferred from glycerol stocks into 2 separate deep well plates containing 1ml of YPD liquid media. One deep well was inoculated using 10μl of inoculum and grown overnight (roughly 23hrs) at 21⁰C, while the other deep well was inoculated using 3ul of inoculum and grown overnight (roughly 16 hrs) at 35⁰C. All plates were grown on rotating drums.

Cells were then measured for OD_600_ values via a Tecan Infinite M200 Pro plate reader. 35⁰C cells were diluted down to an OD_600_ value of roughly 0.04 in 300μl YPD liquid media and then grown for another 4-4.5 hours at 35⁰C. 21⁰C cells were diluted down to an OD_600_ value of roughly 0.08 in 300μl YPD liquid media and then grown for another 4-4.5 hours at 21⁰C. All cells were grown on rotating drums. All following pipetting actions were conducted via a Tecan EVO liquid handling robot. Cells were fixed with fresh Paraformaldehyde (4% final concentration) at room temperature for 30 minutes. Cells were centrifuged at 4000 rpm for 60 seconds and supernatant was removed. Cells were then washed 3 times with PBS, with a spin down at 4000 rpm for 60 seconds and removal of supernatant in between each wash. Cells were then incubated with zymolyase 100T (freshly prepared 50μg/mL) for 2.5 minutes, and then quickly centrifuged for 60 seconds at 2000 rpm, supernatant was then removed. Cells were washed 3 times with PBS, with a spin down at 2000 rpm for 60 seconds and removal of supernatant in between each wash. Following zymolyase digestion and subsequent washes, cells were incubated with PBS containing 10μg/mL Hoechst and 4μM FM® 4-64FX for 60 minutes. Cells were centrifuged for 60 seconds at 2000 rpm and supernatant was removed. Cells were then washed 3 times with PBS, with a spin down at 2000 rpm for 60 seconds and removal of supernatant in between each wash.

Cells were kept at 4°C in the dark for one night. The following day, cells were measured for OD_600_ readings and diluted to an OD_600_ value of 0.02. Cells were then transferred to 384 well PDL coated CellCarrier Ultra imaging plates (PerkinElmer). Quadrants 1 and 2 were reserved for 35°C cells and quadrants 3 and 4 for 21°C cells.

Cells were imaged with an Opera Phenix high content imager (PerkinElmer) using a 63x water objective (C-apochromat, NA 1.15, WD 0.6 mm). Images for GFP fluorescence (Ex 488/Em522), Hoechst staining (Ex 375/Em456) and FM® 4-64FX staining (Ex 488/Em 706) were acquired from 4 fields in each well in triplicate.

#### Quantification of GFP signal

GFP signal was assessed using custom plugins written for the open-source software, ImageJ (NIH, Bethesda, MD). These plugins are publicly available at research.stowers.org/imagejplugins. Cellular compartments were defined as plasma membrane, nuclear membrane, and nucleus. These compartments were experimentally defined by staining with FM4-64 and Hoechst. Segmentation was performed by first segmenting the plasma membrane enclosed space.

Membrane segmentation proceeded as follows. First, FM4-64 images were 2 x 2 boxcar smoothed. Next, a Sobel filter was applied and then thresholded at 5% of its maximum intensity to create a membrane mask. After clearing objects touching the image edges, a binary close operation was performed twice to close membrane gaps. This was followed by a skeletonize operation to thin membranes to a single pixel and then dilated once. These were then filled to create cell masks. Membrane skeletons were removed from these cell masks to ensure separation between cells. Finally, cells were filtered to be no smaller than 20 pixels and no larger than 700 pixels (pixel size is 0.190 microns) and cells were dilated once while preventing merging to retain original boundaries.

Once the cell boundaries were defined, the nucleus was segmented as follows. Firstly, Hoechst images were Gaussian blurred with a standard deviation of 1 pixel. Then each pixel greater than 50% of the difference between the maximum and minimum Hoechst signal in each cell was counted as nuclear. Plasma membrane and nuclear membrane compartments were defined by eroding each compartment once and then dilating twice to obtain the boundary.

Once compartments were identified, average intensities and areas were calculated for each compartment. In addition, the intensity standard deviation was calculated for each cell. Analysis was performed using a multithreaded algorithm where each thread operates on a single image.

Data were then aggregated as follows. Firstly, the average cellular intensity (autofluorescence) of a well containing cells without GFP expression was measured at 21°C and 35°C. Those average intensities were subtracted from all cells at the equivalent temperatures. Cells were further filtered on the ratio of the nuclear area to the cellular area with allowed values between 5% and 40%. Next averages and standard errors for all parameters across all of the cells each well were calculated. Finally, derivative parameters were calculated from average measurements including the nuclear ratio (nuclear intensity/whole cell intensity), membrane ratio (plasma membrane intensity/cytoplasmic intensity), and nuclear membrane ratio (nuclear membrane intensity/nuclear intensity). Standard errors were propagated through the ratio calculations by monte carlo methods assuming Gaussian error distributions for the underlying parameters and 100 simulations. Finally, differences in these ratios as well as the overall intensity and puncta area were calculated between 21°C and 35°C.

Metrics were plugged into a Spotfire™ along with an interactive image viewer showing representative side-by-side images from the different temperatures. This was used to manually validate strains showing strong differences in selected categories.

#### Gene Set Enrichment Analysis of proteins with changes in abundance

We compared the median fluorescence intensity of the 21°C group and 35°C group. The *Z* score was calculated based on the medians of the 35°C and 21°C samples and the median absolute deviation (MAD) of the 21°C sample ((median_35_ -median_21_)/MAD_con_). *Z* scores of -1.65 and 1.65 representing two MADs from the median of 21°C were chosen as cutoff values.

An abundance change profile was created so that each gene in the GFP collection was associated with a Z score as calculated above. The profiles were analyzed by GSEA in pre-rank mode and “c5.bp.v6.2.symbols.gmt” gene sets. All default parameters were used except that the minimum and maximum gene set sizes were set to 5 and 1000, respectively. Enrichment maps were generated with the Enrichment Map Visualization plugin developed for Cytoscape. All default parameters were used except that p-value cut-off were set to 0.05, FDR cut-off of 0.25, and Jacard coefficient cutoff of 0.25.

#### Quantification of protein localization changes using DeepLoc

The published DeepLoc package was used to classify the localization of each proteins at both temperatures. The method can be broken down into data preparation, transfer learning, model evaluation, and finally applying the model to identify significant localization for each protein.

##### Data Preparation

To extract single cell images for DeepLoc model transfer learning, we selected the strains that only show single localization class based on the YEAST GFP FUSION LOCALIZATION DATABASE (https://yeastgfp.yeastgenome.org/, Table single_class_ORFv2.csv) For each strain, we generated single cell segmentation mask based on bright field images using the pre-trained mask-rcnn model (https://github.com/alexxijielu/yeast_segmentation). Then we generated CellProfiler protocol (Kai-yeast-v6.cpproj) (McQuin et al., 2018) to get single cell coordinates, GFP signal intensity and shape from each image. To match the single cell crop dimension (64×64) in DeepLoc, we resized our image sets by 150%, using the Inter-linear interpolation. Single cell images were cropped from GFP channel and the matching nucleus channel. We manually selected single cell images for 17 classifications: actin, bud, bud_neck, cell_periphery, cytoplasm, Endosome, ER, Golgi, mitochondria, none, nuclear_periphery, nucleolus, nucleus, peroxisome, spindle_pole, vacuolar_membrane, vacuole. From each class, we randomly picked 1137 single cells to form our dataset (19329 cells). This dataset was further divided into training (80%) and test (20%) sets.

##### Transfer Learning

Our transfer leaning protocol follows the one described by Kraus, et al. 2017(Kraus et al., 2017) with some modifications. In brief, we initialized a new network loaded with parameters that was trained from the Chong et all 2005 cell dataset for every layer except the final layer. We added a dropout layer to the last fully connected layer (fc2) to prevent overfitting. We transferred and trained the network by feeding in increasing numbers of our training images per class (0, 1, 3, 5, 10, 25, 50, 100, 250, 500 and 1000 samples from each class). For each of these dataset size, five distinct training sets were generated by random sampling from the entire training set. All the parameters were optimized with the Adam optimization, using a learning rate of 0.003 for at least 500 iterations and batch size of 128. To further visualize the separation of 17 classes, we plotted the t-SNE map to visualize the activations on the last layer before the final layer -- fc2_drop layer, for 3866 single cells in our test dataset. We used the Sklearn TSNE implementation to plot 512 features of 3866 cells, with PCA embedding, perplexity of 62 and 5000 iterations. As the numbers of training images per class increases, the separation of 17 classes on t-SNE map becomes more visible.

##### Model Evaluation

We conclude that feeding in 1000 training samples per class produced a model with highest accuracy (0.83) in our test datasets. To further evaluate this model, we predicted 2083 strains in the YEAST GFP FUSION LOCALIZATION DATABASE that show single localization. For each strain we selected cells with GFP signal falls between 0.75-0.9 quantile of all the cells from that strain, and calculated the mean prediction value of each class. Strains with lower than 20 cells predicted were excluded. For each strain the localization class that gets the highest mean prediction value was set as the predicted localization of that strain. We compare the predicted localization and the true localization (based on the YEAST GFP FUSION LOCALIZATION DATABASE) to form the confusion matrix.

The confusion matrix gave 0.89 precision and 0.81 recall value, comparable to the test datasets. Recall value is defined as that among the strains annotated in a given class, how many are determined by DeepLoc to be the same class.

##### Identifying significant localization and GFP abundance changes at higher temperature

We used the new learned model to predict all the strains grown at 35°C as we did with our datasets at 21°C. For each strain we calculated t-score between 21°C and 35°C using Welch’s t-test for all 16 classes. Next for each localization class, we selected strains with dominant localization in this class at 21°C and calculated the Z-score for each strain based on the distribution of their t-score, which represents the localization difference. We applied the same statistic method to the GFP intensity as well. Strains with the absolute value of Z-score larger than 1.96 were considered to show significant change in the corresponding localization or GFP intensity. We thus divided the strains into 4 categories: strains with significant change in both GFP intensity and dominant localization, strains with significant change in only the GFP intensity, strains with significant change in only the dominant localization, and strains with no significant change in either GFP intensity or dominant localization.

To visualize significant localization changes globally, we used Cytoscape to generate a network representing strains with t-score larger than 15 in at least one of the localization classes. Strains with more than one significant localization change were considered as moving from one location to the other. We also scored the normalized intensity change by (35C_meanIntensity - 21C_meanIntensity)/21C_meanIntensity and present it as the color of each node of the strain. t-score was encoded into the thickness of the arrow and direction to indicate magnitude and directional change of the localization.

#### Limited Proteolysis and Mass Spectrometry (LiP-MS)

##### Sample Preparation

Three biological replicates of wild type yeast cells (BY4741) were cultured with YEPD either at 21°C or 35°C overnight (∼18hrs) in glass tubes (10mL per tube) on rotating drums to OD_600_ around 2. Pre-warmed/chilled fresh YEPD was used to dilute the cultures, which were grown for an additional 6hrs to OD_600_ around 1. 120mL of cell culture for each replicate were briefly centrifuged at 3,000 g to collect the cells. Cells were washed once with cold lysis buffer (50mM HEPES, pH 7.5, 150 mM NaCl, 1mM MgCl_2_, 1% Triton X100) and resuspended in 1mL lysis buffer. The cell suspensions were flash frozen as droplets (∼3mm) in pre-chilled 50 mL grinding jars in liquid nitrogen and grinded for 3min at 30 Hz in a mixer mill (Retsch MM400). After melting the samples, cell lysates were transferred to 2mL centrifuge tubes and mixed with an additional 800μl of lysis buffer, which was used to rinse the grinding jar. Cell lysates were clarified by a 10min centrifugation at 16,000 g at 4°C. The concentration of clarified lysate was determined with BCA assay and diluted to 2 mg/mL with buffer (20mM HEPES pH 7.5, NaCl 150mM, 10mM MgCl_2_).

##### Protein Digestion

0.9mL of these diluted lysates were digested with proteinase K at 1:100 (w/w) ratio for 5 min at RT. Another 0.9 mL was saved as control without proteinase K digestion. The proteinase K-digested and control samples were immediately terminated by 7 M guanidine hydrochloride (GdnHCl) and boiled for 5 min. After cooling down to room temperature, samples were reduced with 5 mM TCEP (tris(2-carboxyethyl)phosphine) and alkylated with 10 mM chloroacetamide in dark for 30 min. Samples were diluted with 0.1 M NH_4_HCO_3_ to reduce the concentration of GdnHCl to 0.5 M and digested with trypsin (Promega, 18μg, which is 1:50 w/w) for 2hr at 37⁰C with 1mM CaCl_2_. Additional trypsin (12μg, which is 1:75 w/w) was added to each sample to digest overnight. The samples were next acidified with formic acid (5%) and concentrated using Empore Solid Phase Extraction Cartridges prewashed with 1mL of methanol and 1mL of 0.1% Trifluoroacetic acid (TFA). The peptides were eluted with 80% acetonitrile containing 0.1% TFA after washing the cartridge with 1mL of 0.1% TFA.

##### MS Data Acquisition

All peptide samples were dried using a SpeedVac vacuum concentrator (Thermo Scientific Savant) and resuspended in 150 ml 5% acetonitrile, 0.1% formic acid (Buffer A). Peptides were loaded on a split-triple-phase fused-silica micro-capillary column (Florens and Washburn, 2006) and placed in-line with an Agilent 1260 series quaternary HPLC pump and Velos Pro Orbitrap or Velos Orbitrap Elite mass spectrometers (Thermo Fisher Scientific). A fully automated 12-step MudPIT chromatography run was carried out, as described in (Florens and Washburn, 2006). Each full MS scan (400-1600 m/z) was followed by 15 data-dependent MS/MS scans. MS1 scans were acquired in Orbitrap at 60,000 resolution, while ddMS2 were acquired in the ion trap with Rapid Scan settings. The number of the micro scans was set to 1 for both MS and MS/MS. MS1 AGC target was set to 1.00E+06, MS2 AGC targets at 1.00E+04, and MS2 max injection times at 150 ms. MS1 charge states were between 2-5. MS2 collision energy was set at 35%. Dynamic exclusion settings were: 2 repeat counts; 30 s repeat duration; exclusion list size of 500; and 90 s exclusion duration.

##### MS Data Searching and Processing

Peak files were extracted using RawDistiller (Zhang et al., 2011) and searched using ProLuCID (v. 1.3.3) (Xu et al., 2015) against a database consisting of 5,846 *S. cerevisiae* non-redundant proteins (downloaded from NCBI 11-01-2016), 193 usual contaminants (such as human keratins, IgGs, and proteolytic enzymes), and, to estimate false discovery rates (FDRs), 6,039 randomized amino acid sequences derived from each non-redundant protein entry. All cysteines were considered as fully carboxamidomethylated (+57.0215 Da statically added), while methionine oxidation was searched as a differential modification. Mass tolerances of 7 ppm for precursor and 800 ppm for fragment ions were used. MS/MS datasets generated for trypsin-digested peptides were searched with KR peptide-end requirements, while no enzyme specificity was required for the MS/MS datasets generated for the proteinase K-digested peptides. DTASelect v1.9 (Tabb et al., 2002) and *swallow* v. 0.0.1 (https://github.com/tzw-wen/kite) were used to filter ProLuCID search results at given FDRs at the spectrum, peptide, and protein levels. Here, all controlled FDRs were less than 1%. To estimate relative protein levels and to account for peptides shared between proteins, normalized spectral abundance factors (dNSAFs) were calculated for each detected protein using our in-house developed software, NSAF7 v0.0.1, as described in (Zhang et al., 2010).

##### Post-translational Modification Searches

The MS/MS dataset for the samples only digested with trypsin was further queried for modified peptides in a recursive manner. A total of eight differential modification searches were set up to query for oxidized methionines (+15.9949 Da) in combination with: 1) acetylated lysines, serines, and threonines (+42.0106 Da); 2) formylated arginines and lysines (+27.9949 Da); 3) hydroxylated aspartates, tyrosines, histidines, and prolines (+15.9949); 4) mono- (+14.0157 Da) and 5) dimethylated (+28.0314 Da) arginines and lysines; 6) trimethylated (+42.0470) lysines; 7) phosphorylated (+79.9663 Da) serines, threonines, and tyrosines; and 8) ubiquitinated lysines (+114.0429 Da). Mass tolerances of 7 ppm for precursor and 800 ppm for fragment ions were used. MS/MS datasets generated for trypsin-digested peptides were searched with KR peptide-end requirements. After this round of searches, an in-house software, *sqt-merge*(Zybailov et al., 2005), was used to combine the ProLuCID output files (.sqt format) allowing only the best matches out of the eight differential queries to be ranked first based on cross-correlation scores (XCorr) and for normalized differences between XCorrs (DeltaCn) to be recalculated accordingly. DTASelect v1.9 and *swallow* v. 0.0.1 were used to filter ProLuCID search results at given FDRs at the spectrum, peptide, and protein levels. Here, all controlled FDRs were less than 1%. NSAF7 v0.0.1 was used to generate qualitative and quantitative reports on the detected post-translational modifications.

##### Data Availability

The complete mass spectrometry datasets have been deposited to the MassIVE and Proteome Xchange repositories with the following accessions MSV000086790/PXD023924 for the MudPIT LiP-MS analyses and MSV000086791/PXD023925 for the post-translational modifications searches. They may be accessed via FTP using the MSV accessions as username (e.g., ftp://MSV000086791@massive.ucsd.edu) and “winter” as password.

#### 1D profiling method

The distributed spectral count (dSpC) of peptides was used to represent the abundance of each peptide identified. Peptides in both proteinase K and control digestions were analyzed to remove the non-tryptic peptides found in the control samples. These non-tryptic peptides probably arose from the rare non-specific digestion of trypsin. The proteinase K cutting sites were extracted from the remaining non-tryptic peptides and the N-terminal residues of the fragmentation sites were used to represent the recognition sites of proteinase K (PK residues). The dSpC of the non-tryptic peptides bearing such residues was assigned to the extracted PK residues. The dSpC of the same PK residues from different non-tryptic peptides were summed and normalized to the total dSpC of PK residues from the same protein to represent the accessibility of the residue by proteinase K. Normalized dSpC (n-dSpC) of each PK residues from the same protein was plotted against its primary sequence to generate the 1D profile of each protein in Figure 3. To estimate the reproducibility between biological replicates at 21°C, the PK profiles of the same protein from different biological repeats were plotted together and the differential score was calculated by summing the absolute value of the n-dSpC difference for all PK residues between biological repeats. This gave rise to a range of differential score between 0 and 200, with 0 indicates complete overlapping and no difference between the PK profiles from two repeats while the 200 represents the proteins that has no overlap between 1D profiles (21°C vs 21°C, Figure S3B, S3E-G). The same strategy was applied to quantify the differential score of each protein expressed at different temperatures (21⁰C vs 35⁰C). To search for temperature-dependent 1D profile changes, the differential scores of 21⁰C vs 35⁰C (9 pairs) of each protein that was significantly different from its differential scores of 21⁰C vs 21⁰C (3 pairs) across multiple repeats were calculated using one-way ANOVA test (Figure 3C). Additional 2-fold cutoff was applied to select the hits with large changes in 1D profile between two temperatures.

#### 3D profiling method

The available structures for the proteins identified by proteomics were pooled from Uniprot and the corresponding PDB code and chain identifier for each protein were downloaded and used to fetch the PDB files from protein databank (ftp://ftp.wwpdb.org). For proteins that are subunits of a large complex (e.g. ribosome), the structure of the protein itself was extracted and analyzed independently from the parental complex structure. The depth of each atom in a given protein structure was defined as the distance of this atoms to the solvent accessible surface calculated by Michel Sanner’s MSMS program, which generate the triangulated representation of the solvent-excluded surface (Sanner et al., 1996). The default parameter in this program uses probe size of 1.4 Å, which represents the surface that accessible by water molecules. This leads to a surface with multiple tunneled surfaces through the proteins. As the probing molecule we used in this assay is proteinase K, whose active site and substrate binding pocket are much larger than 15 Å, we modified the probe size for the MSMS program to assess the accessibility of residues to proteinase K active site. After testing a range of probe sizes, ranged from 3 to 15 Å, we found that a probe size around 3 Å still generate tunneled surface in the structures and a probe size of 15 Å failed to cover the entire protein with surface triangles. We chose a probe size of 5 Å for this analysis. Using the Bio.PDB package form Biopython (Cock et al., 2009), we calculated the depth of each atom in a given protein. As the proteolytic cleavage occurs on the peptide bond residues, we used the alpha carbon of the N-terminal residue adjacent to the cleavage site to represent the cutting depth. Many PDB files from protein data bank contain errors in the annotation of amino acid position in the primary sequence, therefore, instead of matching the position of PK residues in reference primary sequence to the position of residues in structure, a pairwise alignment algorithm was used to directly map the cutting sites of each peptides on the reference sequence used by the protein structure. To search for the temperature-dependent PK cutting depth changes, the depths of individual cutting sites from three repeats of each temperature were pool and compared with one-way ANOVA test. To strengthen the analysis, we filtered out the proteins whose deposited structures cover less than 80% of the protein primary sequence. For proteins with multiple structures, we manually checked to confirm that most of the structures agree with each other for each individual protein and selected a representative structure to display. The mean cutting depth of proteins produced in each temperature was calculated by averaging the aggregated cutting depth with different dSpC.

#### FastPP for measuring thermostability

Cells were cultured with YEPD at 21⁰C and 35⁰C overnight (∼18hrs) in glass tubes (10mL per tube) on rotating drums to OD_600_ around 1. Pre-warmed/chilled fresh YEPD was used to dilute the culture and grow for additional 4hrs to OD_600_ around 1. Cells were harvested by brief centrifugation at 3,000 g and washed once with reducing buffer (Tris-H_2_SO_4_, 0.1M, pH 9.5). After resuspending in 1mL reducing buffer, 10mM DTT was added and incubated at room temperature for 5min. Cells were washed once with digestion buffer (1.2M sorbitol, 20mM PO_4_, pH 7.4) and resuspended in 500 μl of digestion buffer. Zymolyase was avoided in this assay due to the protease activity in zymolyase causing depletion of plasma membrane version of Fet3p. Freshly prepared 50mg/mL Lallzyme mmx (Lodi Wine Labs) supplemented with 20mg/mL BSA, protease inhibitor cocktail and PMSF was used to digest the cell wall for 5min at room temperature. The digested cells were washed twice with digestion buffer to remove enzymes and chilled down in ice before lysing with 500 μl of lysis buffer (50mM Tris pH7.4, 150mM NaCl, 0.34 mM n-Dodecyl-beta-D-maltopyranoside (DDM), 5% glycerol, plus protease inhibitor cocktail, and PMSF). Cell lysate was clarified by centrifugation at 15,000 g for 10min at 4⁰C. The protein concentration of the lysate was determined by BCA assay. Protein lysate was supplemented with 5mM DTT and 10mM CaCl_2_ and then 20 ug protein lysate was aliquoted into each PCR tube. A final reaction volume of 40μL was used for all proteins. Thermolysin was mixed with the samples on ice to a final concentration of 0.15mg/mL and a gradient temperature in a thermocycler was used to heat the samples at specific temperatures for 6 secs. Control samples mixed with thermolysin and kept on ice for the same amount of time was used to normalize the samples digested at elevated temperatures. This basic protocol was used for the following derivations: (1) To evaluate the effect of glycosylation, samples were treated with 15,000 units of Endo H for 30min at room temperature before mixing with thermolysin. (2) To investigate whether the thermostability depends on interaction partners, antiflag resin (A2220 sigma, 100uL resin added to 2mL protein lysate) was used to purify the Fet3-3xflag and the beads were washed with lysis buffer. The thermolysin digestion was performed with bead-anchored Fet3-3xflag. As it is impossible to estimate the amount of proteins on beads, less thermolysin (0.1mg/mL) was used to digest the proteins. For Erg6-3xFlag, which was a more thermostable protein, 25μg protein lysate was added to each tube and 0.3mg/mL thermolysin was used to digest the protein as it unfolded.

#### Protease protection assay

Cells were cultured with YEPD at 21°C and 35°C overnight (∼18hrs) in glass tubes (10mL per tube) on rotating drums to OD_600_ around 1. Pre-warmed/chilled Fresh YEPD was used to dilute the culture and grow for additional 4hrs to OD_600_ around 1. Culture was aliquoted into fractions with the same number of cells for each treatment. 1×10^8^ Cells were washed and incubated for 20min with cold YEPD containing 20mM potassium fluoride (KF) and sodium azide (NaN_3_) on ice. Cells were either directly lysed with 5min treatment of 0.1 M NaOH at room temperature according to a previous protocol or digested with the Lallzyme method as above. To quantify the distribution of Fet3p between plasma membrane and endoplasmic reticulum (both nuclear and cortical ER), the Lallzyme-digested cells were resuspended in 400μl of DB buffer (1.4M sorbitol, 25mM Tris pH 7.5, 10mM NaN3, 10mM KF, 2mM MgCl_2_) and mixed with 100μl of pronase (2500U/mL in DB buffer) or 100μl of DB buffer as control for 30min at RT. Both samples were washed three times with DB buffer containing protease inhibitor cocktail and PMSF to remove pronase. Cells were then lysed with sample buffer for western blot.

#### Mass spectrometry to identify interaction partners of Fet3p

##### Sample Preparation

Cells expressing 3xflag and 6xhis-tagged Fet3p were cultured with YEPD at 21⁰C and 35⁰C overnight (∼18hrs) in glass tubes (10mL per tube) on rotating drums to OD_600_ around 1. Pre-warmed/chilled fresh YEPD was used to dilute the culture and grow for an additional 4hrs to OD_600_ around 1. About 120 OD_600_ cells from each culture were digested with the Lallzyme protocol as above and lysed with lysis buffer (50 mM Tris, pH 7.4, with 150 mM NaCl, 0.5%triton, protease inhibitor cocktail and 1mM PMSF, 5%Glycerol). Lysate was then clarified with 30,000g centrifugation at 4°C and the soup was incubated with 100μl of antiflag beads for 1hr at 4°C. The beads were washed with 30 volumes of cold TBS pH 7.5 to remove nonspecific binding proteins. The proteins were eluted with Glycine/HCl buffer, pH 3.3 from beads and neutralized with Tris buffer to pH 7.4. This particular Mass spectrometry assay was done by the proteomics center at the Buck Institute:

##### Chemicals

HPLC-grade acetonitrile and water were obtained from Burdick & Jackson (Muskegon, MI). Reagents for protein chemistry, including iodoacetamide, dithiothreitol (DTT), ammonium bicarbonate, and formic acid (FA) were purchased from Sigma-Aldrich (St. Louis, MO). Sequencing-grade trypsin was purchased from Promega (Madison, WI).

##### Proteomic sample preparation

Enriched yeast samples from an anti-flag pulldown experiment, containing 2-4 µg protein, were concentrated from 400 µL to 25 µL using 0.5 mL centrifugal filters, after which the concentrated samples were loaded on a 1D SDS-PAGE system. The gel bands were diced and collected in tubes, and vortexed several times in dehydration buffer (25 mM ammonium bicarbonate in 50% acetonitrile and water). The gel samples were concentrated to dryness (speedvac) and reduced with 10 mM dithiothreitol for 1 hour at 56°C with agitation. Samples were subsequently alkylated with 55 mM iodoacetamide for 45 minutes at room temperature in the dark. The diced gels pieces were washed with 25 mM ammonium bicarbonate in water, and then dehydrated once again with the dehydration buffer. The samples were then concentrated to dryness (speedvac) after which they were incubated with 250 ng trypsin for 30 minutes at 4°C, and then digested overnight at 37°C with agitation. The following morning, the digestions were subjected to water and then 50% acetonitrile and 5% formic acid in water. After each addition of a solution, the aqueous digest in each sample was collected into a new tube. These pooled peptide extractions were concentrated for two hours to reach dryness (speedvac), and then re-suspended in 0.2% formic acid. The re-suspended peptide samples were desalted with Zip Tips containing a C18 disk, concentrated (speedvac) and re-suspended in a solution containing “Hyper Reaction Monitoring” (Biognosys) indexed retention time peptide standards (iRT) and 0.2% formic acid in water.

##### Mass Spectrometry System

Briefly, samples were analyzed by reverse-phase HPLC-ESI-MS/MS using an Eksigent Ultra Plus nano-LC 2D HPLC system (Dublin, CA) with a cHiPLC system (Eksigent) which was directly connected to a quadrupole time-of-flight (QqTOF) TripleTOF 6600 mass spectrometer (SCIEX, Concord, CAN). After injection, peptide mixtures were loaded onto a C18 pre-column chip (200 µm x 0.4 mm ChromXP C18-CL chip, 3 µm, 120 Å, SCIEX) and washed at 2 µl/min for 10 min with the loading solvent (H_2_O/0.1% formic acid) for desalting. Subsequently, peptides were transferred to the 75 µm x 15 cm ChromXP C18-CL chip, 3 µm, 120 Å, (SCIEX), and eluted at a flow rate of 300 nL/min with a 3 hr gradient using aqueous and acetonitrile solvent buffers.

##### Data-dependent acquisitions

For peptide and protein identifications the mass spectrometer was operated in data-dependent acquisition (DDA) mode, where the 30 most abundant precursor ions from the survey MS1 scan (250 msec) were isolated at 1 m/z resolution for collision induced dissociation tandem mass spectrometry (CID-MS/MS, 100 msec per MS/MS, ‘high sensitivity’ product ion scan mode) using the Analyst 1.7 (build 96) software with a total cycle time of 3.3 sec as previously described (Christensen et al., 2018).

##### Mass-spectrometric data processing and bioinformatics

Mass spectrometric data-dependent acquisitions (DDA) were analyzed using the database search engine ProteinPilot 5.0.0.0, 4769 (SCIEX) using the Paragon algorithm (5.0.0.0, 4767) (Shilov et al., 2007). The search parameters were set as follows: trypsin digestion, cysteine alkylation set to iodoacetamide, acetylation emphasis, and species *Saccharomyces cerevisiae* (with 6723 proteins in the database). Trypsin specificity was assumed as C-terminal cleavage at lysine and arginine. Processing parameters were set to “Biological modification” and a thorough ID search effort was used. For database searches, a cut-off peptide confidence value of 99 was chosen. The Protein Pilot false discovery rate (FDR) analysis tool, the Proteomics System Performance Evaluation Pipeline (PSPEP algorithm) (Shilov et al., 2007) provided a global FDR of 1% and a local FDR at 1% in all cases.

##### Data availability

The mass spectrometric raw data are deposited at ftp://MSV000086243@massive.ucsd.edu with the MassIVE ID MSV000086243 (password: winter) (upon removal of the password data will be available at ftp://massive.ucsd.edu/MSV000086243/) it is also available at ProteomeXchange with the ID PXD021862.

#### Co-immunoprecipitation

Cells expressing Fet3-3xFlag-URA and a target protein tagged with GFP-HIS were cultured with YEPD at 21⁰C and 35⁰C overnight (∼18hrs) in glass tubes (10mL per tube) on rotating drums to OD_600_ around 1. Pre-warmed/chilled YEPD was used to dilute the culture and grow for an additional 4hrs to OD_600_ around 1. About 20 OD_600_ cells from each culture were reduced with 1mM DTT for 5min then digested with 500 µL of 50mg/mL Lallzyme (protocol as above) and lysed with 1mL lysis buffer (50 mM Tris, pH 7.4, with 150 mM NaCl, 0.5%triton, protease inhibitor cocktail and 1mM PMSF, 5% Glycerol). Lysate was then clarified with 15,000g centrifugation at 4°C for 10min and the soup was incubated with 40ul of anti-flag beads (A2220, Sigma) for 1hr at 4°C. The beads were washed with 20 times beads volume of chilled (4°C) TBS pH 7.6 to remove nonspecific binding proteins. Finally, the beads were resuspended in 50uL of 1x sample buffer and boiled for 5mins to elute the proteins from the beads. 10uL of eluate was loaded onto an SDS-PAGE gel and ran for 1hr at 120V, then transferred to 0.45μm PVDF membrane at 350mAmp for 80min then blocked in 5% milk in TBST and then probed with anti-flag and anti-GFP antibodies (Anti-GFP, SAB4301138) For Fet3-3xFlag, Elo3-GFP, a different lysis protocol was used due to Elo3-GFP being quickly degraded and otherwise lost unless the vacuole was first separated from the protein lysate. Cells were grown and harvested as described above but after spheroplasting with Lallzyme, the cells were resuspended in 250 µL homogenization buffer (0.6M Sorbitol, 50mM Tris pH 7.4, 150mM NaCl, 5% Glycerol, and PIs and PMSF). The cells were then homogenized for 90 sec to break them apart and release the vacuoles then clarified with 15,000g centrifugation for 5min at 4°C. The soup was discarded, and the pellet was resuspended in 750 µL of the triton lysis buffer described above and then 40 µL of anti-flag beads were added.

#### Lipid profiling

Three replicates of wild type and *Δfet3* cells were cultured with SC complete medium at 21⁰C and 35⁰C overnight (∼18hrs) in glass tubes (10mL per tube) on rotating drums to OD_600_ around 1. Pre-warmed/chilled fresh YEPD was used to dilute the culture and grow for additional 4hrs to OD_600_ around 1. 40 OD_600_ cells were collected and flash frozen with liquid nitrogen. Cells were resuspended in 500ul NH_4_HCO_3_ grinded with glass beads. Cell lysate was extracted twice (5min vortex) with 1.5mL of CHCl_3_/Methanol (2:1) contains 200ng/mL FA 17:0 Saturated. The chloroform layer was extracted and combined, dried under nitrogen and reconstituted with chloroform.

The reconstituted lipid extracts were analyzed using Agilent 6520 quadruple time-of-flight mass spectrometer (6520-QToF) coupled to Agilent 1260 Infinity LC system. The HPLC system includes modules, which included degasser (G1322A), binary pump (G1312B), thermo-regulated column compartment (G1330B), and HiPALS autosampler (G1367E). Chromatographic separation of lipids was performed on HALO AQ C18 (2.1×150mm, 2.7µm) column. The solvent system consisted of 20 mM ammonium acetate in 5% acetonitrile containing 0.1% acetic acid (A) and 20 mM ammonium acetate in 95% acetonitrile containing 0.1% acetic acid (B). The starting gradient conditions were 95% B at a flow rate of 0.3 mL/min up to 2 min. The following gradient program was used: 2–22 min, 5%–95% B, 22– 28 min 95% B and 29–35 min 5% B. Samples were kept at 4°C, and the injection volume was 10 μL. High resolution MS1 was performed using 6520 QTOF mass spectrometer fitted with a Dual-Spray Electrospray Ionization (ESI) source. The instrument was operated at a mass resolution of ∼20,000 for TOF MS1 scan using 2GHx extended dynamic range mode. The ionization parameters were set as follows: gas temperature (TEM) 350°C; drying gas, 9 L/min; Vcap, 2,500 V; nebulizer gas, 35 psi; fragmentor, 125 V; and skimmer, 65 V. MS1 acquisition was operated in the negative ion scanning mode for a mass range of 50–1,000 m/z. The HPLC-MS data was acquired using Agilent MassHunter Workstation (B.05.00), Agilent MassHunter Qualitative Analysis Software (B.07.00) and Mass Profiler Professional (B.12.0). Individual lipids were identified by generating Agilent’s Personal Compound Database Library (PCDL) by importing a .csv file from LIPID MAPS Structure Database (LMSD). Samples were normalized according to the total phosphatidylcholines in each sample.

#### Whole genome sequencing

Three replicates of By4741 were cultured at 21°C and 35°C for 20hrs and refreshed for 4hrs. 15 mL mid-log cultures (5 OD_600_ cell in total) were used to purify the genomic DNA using Masterpure DNA purification kit. Libraries were prepared from these genomic DNA samples. For the first set of samples, 500ng was sonicated to 300-bp fragments using a S220 Focused-ultrasonicator (Covaris) by adjusting the treatment time to 50 seconds. For the second set of samples, 500ng was sonicated to 450-bp fragments by adjusting the treatment time to 58 seconds. These conditions were repeated for all samples, and both sets of samples were size selected following sonication using a PippinHT (Sage Science). The 300bp set of samples were size selected from 150-450bp, and 450bp set of samples were size selected from 200-600bp. Post sonication and size selection, samples were combined in order to start with more material for library construction. Libraries were prepared from the pooled sample material using the KAPA HTP Library Prep Kit (Roche, Cat No. KK8234) with 15 cycles of PCR and using 1:50 dilution of NEXTflex DNA barcodes (Perkin Elmer, Cat No. NOVA-514104). The resulting libraries were purified using the Agencourt AMPure XP system (Beckman Coulter) then quantified using a Bioanalyzer (Agilent Technologies) and a Qubit fluorometer (Life Technologies). Libraries were re-quantified, normalized, pooled and sequenced on an Illumina MiSeq instrument as 75-bp paired reads (Illumina, Cat No. MS-102-3001). Following sequencing, Illumina Real Time Analysis version 1.18.54.4 and bcl2fastq2 v2.20 were run to demultiplex reads and generate FASTQ files. Following alignment to sacCer3 with bwa mem, reads were deduplicated and analyzed with GATK 4.1.9 using a pipeline based on GATK Best Practices. Briefly, two rounds of bootstrapping with Base Quality Score Recalibration were applied and variants were called using HaplotypeCaller with a ploidy of 1. Variants were annotated using Ensembl Variant Effect Predictor with *Saccharomyces cerevisiae* R64-1-1. See also Table S5.

Yeast isolates were examined for differences in ploidy between conditions by calculating RPM coverage across the genome for each isolate, and then examining the ratio of the median RPM coverage of 5 kb bins across a chromosome to the median of RPM values for all genomic bins. All chromosomes were present in equal ratios across all strains.

The Table S5 lists 4 classes of variants: SNPs that are unique to at least one isolate, or which were not able to be called across all isolates (unique_snp), SNPs or INDELs that are identical across all six isolates (common_snp or common_indel, respectively), thus indicating a uniform difference from S288C, and INDELs that are unique to at least one isolate, or unable to be called across all six (unique_indel). The genotype of each isolate is indicated in the table by an integer indicating which alternative allele is present. A value of 0 indicates a match to the S288C reference, and a value of “.” indicates no call was made for that isolate at that position.

### QUANTIFICATION AND STATISTICAL ANALYSIS

All experiments were repeated multiple times to confirm reproducibility. Data, except for the GFP screen, are representative of at least two independent repeats. All quantifications are presented as the means ± standard error of mean (SEM). Statistical test for each bar graph in figures was determined by ANOVA or unpaired t test with Welch’s correction. n.s. or ns, not significant; *p < 0.05; **p <0.01; ***p < 0.001.

## Supplemental tables

Table S1: Quantification of GFP intensity of different strains and their subcellular localization from the GFP screen.

Table S2: Protein- and peptide-level qualitative and quantitative information for all proteins detected by MudPIT analyses of whole cell extracts from *Saccharomyces cerevisiae* grown at 21°C and 35°C.

Table S3: Hits from the 1D profile analysis and the 3D profiling analysis of the hits with solved reference structures.

Table S4: Hits from the 3D profile analysis and their peptides with or without PTMs.

Table S5: SNPs and genetic variations identified in whole genome sequencing.

Table S6: Temperature-specific protein-protein interaction identified by mass spectrometry and the physical interacting partners of Fet3p identified in previous large-scale screens.

## Supplemental movies

We recommend movie S1 to be viewed using QuickTime player.

**Movie S1**. Map of cutting sites in 3D structure for NP_011651, NP_011467, NP_014769, NP_013127, and NP_013774. Cutting sites from all biological repeats of each temperature were pooled. The scale of ball size, which reflects the frequency of proteinase K cutting, is annotated in the right corner of the structure. Illustration was modified from 1fnt (Whitby et al., 2000), 3jcp (Luan et al., 2016), 3mil (Ma et al., 2011), 1yaa (Jeffery et al., 1998), and 1zpu (Taylor et al., 2005), respectively.

## References

Ashburner, M., Ball, C.A., Blake, J.A., Botstein, D., Butler, H., Cherry, J.M., Davis, A.P., Dolinski, K., Dwight, S.S., Eppig, J.T., et al. (2000). Gene Ontology: tool for the unification of biology. Nat. Genet. 25, 25–29.

Bosson, R., Jaquenoud, M., and Conzelmann, A. (2006). GUP1 of Saccharomyces cerevisiae encodes an O-acyltransferase involved in remodeling of the GPI anchor. Mol. Biol. Cell 17, 2636–2645.

Cappelletti, V., Hauser, T., Piazza, I., Pepelnjak, M., Malinovska, L., Fuhrer, T., Li, Y., Dörig, C., Boersema, P., Gillet, L., et al. (2021). Dynamic 3D proteomes reveal protein functional alterations at high resolution in situ. Cell 184, 545–559.e22.

Christensen, D.G., Meyer, J.G., Baumgartner, J.T., D’Souza, A.K., Nelson, W.C., Payne, S.H., Kuhn, M.L., Schilling, B., and Wolfe, A.J. (2018). Identification of Novel Protein Lysine Acetyltransferases in Escherichia coli. MBio 9.

Cock, P.J.A., Antao, T., Chang, J.T., Chapman, B.A., Cox, C.J., Dalke, A., Friedberg, I., Hamelryck, T., Kauff, F., Wilczynski, B., et al. (2009). Biopython: freely available Python tools for computational molecular biology and bioinformatics. Bioinformatics 25, 1422–1423.

De Silva, D.M., Askwith, C.C., Eide, D., and Kaplan, J. (1995). The FET3 gene product required for high affinity iron transport in yeast is a cell surface ferroxidase. J. Biol. Chem. 270, 1098–1101.

Dishman, A.F., and Volkman, B.F. (2018). Unfolding the mysteries of protein metamorphosis. ACS Chem. Biol. 13, 1438–1446.

Feng, Y., De Franceschi, G., Kahraman, A., Soste, M., Melnik, A., Boersema, P.J., de Laureto, P.P., Nikolaev, Y., Oliveira, A.P., and Picotti, P. (2014). Global analysis of protein structural changes in complex proteomes. Nat. Biotechnol. 32, 1036–1044.

Florens, L., and Washburn, M.P. (2006). Proteomic analysis by multidimensional protein identification technology. Methods Mol. Biol. 328, 159–175.

Fontana, A., de Laureto, P.P., de Filippis, V., Scaramella, E., and Zambonin, M. (1999). Limited proteolysis in the study of protein conformation. In Proteolytic Enzymes: Tools and Targets, E.E. Sterchi, and W. Stöcker, eds. (Berlin, Heidelberg: Springer Berlin Heidelberg), pp. 253–280.

Fontana, A., de Laureto, P.P., Spolaore, B., Frare, E., Picotti, P., and Zambonin, M. (2004). Probing protein structure by limited proteolysis. Acta Biochim. Pol. 51, 299–321.

Funakoshi, M., and Hochstrasser, M. (2009). Small epitope-linker modules for PCR-based C-terminal tagging in Saccharomyces cerevisiae. Yeast 26, 185–192.

Gasch, A.P., Spellman, P.T., Kao, C.M., Carmel-Harel, O., Eisen, M.B., Storz, G., Botstein, D., and Brown, P.O. (2000). Genomic expression programs in the response of yeast cells to environmental changes. Mol. Biol. Cell 11, 4241–4257.

Gene Ontology Consortium (2021). The Gene Ontology resource: enriching a GOld mine. Nucleic Acids Res. 49, D325–D334.

Giaever, G., Chu, A.M., Ni, L., Connelly, C., Riles, L., Véronneau, S., Dow, S., Lucau-Danila, A., Anderson, K., André, B., et al. (2002). Functional profiling of the Saccharomyces cerevisiae genome. Nature 418, 387–391.

Hartl, F.U., Bracher, A., and Hayer-Hartl, M. (2011). Molecular chaperones in protein folding and proteostasis. Nature 475, 324–332.

Henderson, C.M., Zeno, W.F., Lerno, L.A., Longo, M.L., and Block, D.E. (2013). Fermentation temperature modulates phosphatidylethanolamine and phosphatidylinositol levels in the cell membrane of Saccharomyces cerevisiae. Appl. Environ. Microbiol. 79, 5345–5356.

Hopper, A.K. (2013). Transfer RNA post-transcriptional processing, turnover, and subcellular dynamics in the yeast Saccharomyces cerevisiae. Genetics 194, 43–67.

Huh, W.-K., Falvo, J.V., Gerke, L.C., Carroll, A.S., Howson, R.W., Weissman, J.S., and O’Shea, E.K. (2003). Global analysis of protein localization in budding yeast. Nature 425, 686–691.

Jarosz, D.F., and Lindquist, S. (2010). Hsp90 and environmental stress transform the adaptive value of natural genetic variation. Science 330, 1820–1824.

Jeffery, C.J., Barry, T., Doonan, S., Petsko, G.A., and Ringe, D. (1998). Crystal structure of Saccharomyces cerevisiae cytosolic aspartate aminotransferase. Protein Sci. 7, 1380–1387.

Jin, L., Baker, B., Mealer, R., Cohen, L., Pieribone, V., Pralle, A., and Hughes, T. (2011). Random insertion of split-cans of the fluorescent protein venus into Shaker channels yields voltage sensitive probes with improved membrane localization in mammalian cells. J. Neurosci. Methods 199, 1–9.

Jones, D.T., and Cozzetto, D. (2015). DISOPRED3: precise disordered region predictions with annotated protein-binding activity. Bioinformatics 31, 857–863.

Kimchi-Sarfaty, C., Oh, J.M., Kim, I.-W., Sauna, Z.E., Calcagno, A.M., Ambudkar, S.V., and Gottesman, M.M. (2007). A “silent” polymorphism in the MDR1 gene changes substrate specificity. Science 315, 525–528.

Kisser, B., and Goodwin, H.T. (2012). Hibernation and overwinter body temperatures in free-ranging thirteen-lined ground squirrels, *Ictidomys tridecemlineatus*. Am. Midl. Nat. 167, 396–409.

Klausner, R.D., and Rouault, T.A. (1993). A double life: cytosolic aconitase as a regulatory RNA binding protein. Mol. Biol. Cell 4, 1–5.

Kramer, G., Shiber, A., and Bukau, B. (2019). Mechanisms of cotranslational maturation of newly synthesized proteins. Annu. Rev. Biochem. 88, 337–364.

Kraus, O.Z., Grys, B.T., Ba, J., Chong, Y., Frey, B.J., Boone, C., and Andrews, B.J. (2017). Automated analysis of high-content microscopy data with deep learning. Mol. Syst. Biol. 13, 924.

LeCun, Y., Bengio, Y., and Hinton, G. (2015). Deep learning. Nature 521, 436–444.

Leuenberger, P., Ganscha, S., Kahraman, A., Cappelletti, V., Boersema, P.J., von Mering, C., Claassen, M., and Picotti, P. (2017). Cell-wide analysis of protein thermal unfolding reveals determinants of thermostability. Science 355.

Liu, B., Han, Y., and Qian, S.-B. (2013). Cotranslational response to proteotoxic stress by elongation pausing of ribosomes. Mol. Cell 49, 453–463.

Longtine, M.S., McKenzie, A., Demarini, D.J., Shah, N.G., Wach, A., Brachat, A., Philippsen, P., and Pringle, J.R. (1998). Additional modules for versatile and economical PCR-based gene deletion and modification in Saccharomyces cerevisiae. Yeast 14, 953–961.

Luan, B., Huang, X., Wu, J., Mei, Z., Wang, Y., Xue, X., Yan, C., Wang, J., Finley, D.J., Shi, Y., et al. (2016). Structure of an endogenous yeast 26S proteasome reveals two major conformational states. Proc. Natl. Acad. Sci. USA 113, 2642–2647.

Ma, J., Lu, Q., Yuan, Y., Ge, H., Li, K., Zhao, W., Gao, Y., Niu, L., and Teng, M. (2011). Crystal structure of isoamyl acetate-hydrolyzing esterase from Saccharomyces cerevisiae reveals a novel active site architecture and the basis of substrate specificity. Proteins 79, 662–668.

Maaten, L. van der, and Hinton, G. (2008). Visualizing Data using t-SNE. Journal of Machine Learning Research.

Mayor, S., and Riezman, H. (2004). Sorting GPI-anchored proteins. Nat. Rev. Mol. Cell Biol. 5, 110– 120.

McQuin, C., Goodman, A., Chernyshev, V., Kamentsky, L., Cimini, B.A., Karhohs, K.W., Doan, M., Ding, L., Rafelski, S.M., Thirstrup, D., et al. (2018). CellProfiler 3.0: Next-generation image processing for biology. PLoS Biol. 16, e2005970.

Mészáros, B., Erdos, G., and Dosztányi, Z. (2018). IUPred2A: context-dependent prediction of protein disorder as a function of redox state and protein binding. Nucleic Acids Res. 46, W329–W337.

Meurer, M., Duan, Y., Sass, E., Kats, I., Herbst, K., Buchmuller, B.C., Dederer, V., Huber, F., Kirrmaier, D., Štefl, M., et al. (2018). Genome-wide C-SWAT library for high-throughput yeast genome tagging. Nat. Methods 15, 598–600.

Miller, J.P., Lo, R.S., Ben-Hur, A., Desmarais, C., Stagljar, I., Noble, W.S., and Fields, S. (2005). Large-scale identification of yeast integral membrane protein interactions. Proc. Natl. Acad. Sci. USA 102, 12123–12128.

Minde, D.P., Maurice, M.M., and Rüdiger, S.G.D. (2012). Determining biophysical protein stability in lysates by a fast proteolysis assay, FASTpp. PLoS One 7, e46147.

Mogk, A., Bukau, B., and Kampinga, H.H. (2018). Cellular handling of protein aggregates by disaggregation machines. Mol. Cell 69, 214–226.

Montefusco, D.J., Matmati, N., and Hannun, Y.A. (2014). The yeast sphingolipid signaling landscape. Chem Phys Lipids 177, 26–40.

Murzin, A.G. (2008). Biochemistry. Metamorphic proteins. Science 320, 1725–1726.

Neubert, P., Halim, A., Zauser, M., Essig, A., Joshi, H.J., Zatorska, E., Larsen, I.S.B., Loibl, M., Castells-Ballester, J., Aebi, M., et al. (2016). Mapping the O-Mannose Glycoproteome in Saccharomyces cerevisiae. Mol. Cell Proteomics 15, 1323–1337.

O’Brien, E.P., Vendruscolo, M., and Dobson, C.M. (2014). Kinetic modelling indicates that fast-translating codons can coordinate cotranslational protein folding by avoiding misfolded intermediates. Nat. Commun. 5, 2988.

Onuchic, J.N., Luthey-Schulten, Z., and Wolynes, P.G. (1997). Theory of protein folding: the energy landscape perspective. Annu. Rev. Phys. Chem. 48, 545–600.

Parks, L.W., and Casey, W.M. (1995). Physiological implications of sterol biosynthesis in yeast. Annu. Rev. Microbiol. 49, 95–116.

Pechmann, S., and Frydman, J. (2013). Evolutionary conservation of codon optimality reveals hidden signatures of cotranslational folding. Nat. Struct. Mol. Biol. 20, 237–243.

Pechmann, S., Chartron, J.W., and Frydman, J. (2014). Local slowdown of translation by nonoptimal codons promotes nascent-chain recognition by SRP *in vivo*. Nat. Struct. Mol. Biol. 21, 1100–1105.

Piatigorsky, J., O’Brien, W.E., Norman, B.L., Kalumuck, K., Wistow, G.J., Borras, T., Nickerson, J.M., and Wawrousek, E.F. (1988). Gene sharing by delta-crystallin and argininosuccinate lyase. Proc. Natl. Acad. Sci. USA 85, 3479–3483.

Richter, J.D., and Coller, J. (2015). Pausing on polyribosomes: make way for elongation in translational control. Cell 163, 292–300.

Roth, D.M., Hutt, D.M., Tong, J., Bouchecareilh, M., Wang, N., Seeley, T., Dekkers, J.F., Beekman, J.M., Garza, D., Drew, L., et al. (2014). Modulation of the maladaptive stress response to manage diseases of protein folding. PLoS Biol. 12, e1001998.

Ruan, L., Zhou, C., Jin, E., Kucharavy, A., Zhang, Y., Wen, Z., Florens, L., and Li, R. (2017). Cytosolic proteostasis through importing of misfolded proteins into mitochondria. Nature 543, 443–446.

Rutherford, S.L., and Lindquist, S. (1998). Hsp90 as a capacitor for morphological evolution. Nature 396, 336–342.

Sanner, M.F., Olson, A.J., and Spehner, J.-C. (1996). Reduced surface: An efficient way to compute molecular surfaces. Biopolymers 38, 305–320.

Shalgi, R., Hurt, J.A., Krykbaeva, I., Taipale, M., Lindquist, S., and Burge, C.B. (2013). Widespread regulation of translation by elongation pausing in heat shock. Mol. Cell 49, 439–452.

Shannon, P., Markiel, A., Ozier, O., Baliga, N.S., Wang, J.T., Ramage, D., Amin, N., Schwikowski, B., and Ideker, T. (2003). Cytoscape: a software environment for integrated models of biomolecular interaction networks. Genome Res. 13, 2498–2504.

Shiber, A., Döring, K., Friedrich, U., Klann, K., Merker, D., Zedan, M., Tippmann, F., Kramer, G., and Bukau, B. (2018). Cotranslational assembly of protein complexes in eukaryotes revealed by ribosome profiling. Nature 561, 268–272.

Shilov, I.V., Seymour, S.L., Patel, A.A., Loboda, A., Tang, W.H., Keating, S.P., Hunter, C.L., Nuwaysir, L.M., and Schaeffer, D.A. (2007). The Paragon Algorithm, a next generation search engine that uses sequence temperature values and feature probabilities to identify peptides from tandem mass spectra. Mol. Cell Proteomics 6, 1638–1655.

Singh, A., Severance, S., Kaur, N., Wiltsie, W., and Kosman, D.J. (2006). Assembly, activation, and trafficking of the Fet3p.Ftr1p high affinity iron permease complex in Saccharomyces cerevisiae. J. Biol. Chem. 281, 13355–13364.

Stearman, R., Yuan, D.S., Yamaguchi-Iwai, Y., Klausner, R.D., and Dancis, A. (1996). A permease-oxidase complex involved in high-affinity iron uptake in yeast. Science 271, 1552–1557.

Tabb, D.L., McDonald, W.H., and Yates, J.R. (2002). DTASelect and Contrast: tools for assembling and comparing protein identifications from shotgun proteomics. J. Proteome Res. 1, 21–26.

Taylor, A.B., Stoj, C.S., Ziegler, L., Kosman, D.J., and Hart, P.J. (2005). The copper-iron connection in biology: structure of the metallo-oxidase Fet3p. Proc. Natl. Acad. Sci. USA 102, 15459–15464.

Tokuriki, N., and Tawfik, D.S. (2009). Protein dynamism and evolvability. Science 324, 203–207.

Towns, W.L., and Begley, T.J. (2012). Transfer RNA methytransferases and their corresponding modifications in budding yeast and humans: activities, predications, and potential roles in human health. DNA Cell Biol 31, 434–454.

Tuinstra, R.L., Peterson, F.C., Kutlesa, S., Elgin, E.S., Kron, M.A., and Volkman, B.F. (2008). Interconversion between two unrelated protein folds in the lymphotactin native state. Proc. Natl. Acad. Sci. USA 105, 5057–5062.

Wang, X., Venable, J., LaPointe, P., Hutt, D.M., Koulov, A.V., Coppinger, J., Gurkan, C., Kellner, W., Matteson, J., Plutner, H., et al. (2006). Hsp90 cochaperone Aha1 downregulation rescues misfolding of CFTR in cystic fibrosis. Cell 127, 803–815.

Weill, U., Yofe, I., Sass, E., Stynen, B., Davidi, D., Natarajan, J., Ben-Menachem, R., Avihou, Z., Goldman, O., Harpaz, N., et al. (2018). Genome-wide SWAp-Tag yeast libraries for proteome exploration. Nat. Methods 15, 617–622.

Whitby, F.G., Masters, E.I., Kramer, L., Knowlton, J.R., Yao, Y., Wang, C.C., and Hill, C.P. (2000). Structural basis for the activation of 20S proteasomes by 11S regulators. Nature 408, 115–120.

Xu, T., Park, S.K., Venable, J.D., Wohlschlegel, J.A., Diedrich, J.K., Cociorva, D., Lu, B., Liao, L., Hewel, J., Han, X., et al. (2015). ProLuCID: An improved SEQUEST-like algorithm with enhanced sensitivity and specificity. J. Proteomics 129, 16–24.

Yeger-Lotem, E., Riva, L., Su, L.J., Gitler, A.D., Cashikar, A.G., King, O.D., Auluck, P.K., Geddie, M.L., Valastyan, J.S., Karger, D.R., et al. (2009). Bridging high-throughput genetic and transcriptional data reveals cellular responses to alpha-synuclein toxicity. Nat. Genet. 41, 316–323.

Zhang, Y., Wen, Z., Washburn, M.P., and Florens, L. (2010). Refinements to label free proteome quantitation: how to deal with peptides shared by multiple proteins. Anal. Chem. 82, 2272–2281.

Zhang, Y., Wen, Z., Washburn, M.P., and Florens, L. (2011). Improving proteomics mass accuracy by dynamic offline lock mass. Anal. Chem. 83, 9344–9351.

Zheng, X., Krakowiak, J., Patel, N., Beyzavi, A., Ezike, J., Khalil, A.S., and Pincus, D. (2016). Dynamic control of Hsf1 during heat shock by a chaperone switch and phosphorylation. Elife 5.

Zhou, C., Slaughter, B.D., Unruh, J.R., Guo, F., Yu, Z., Mickey, K., Narkar, A., Ross, R.T., McClain, M., and Li, R. (2014). Organelle-based aggregation and retention of damaged proteins in asymmetrically dividing cells. Cell 159, 530–542.

Zielinska, D.F., Gnad, F., Schropp, K., Wiśniewski, J.R., and Mann, M. (2012). Mapping N-glycosylation sites across seven evolutionarily distant species reveals a divergent substrate proteome despite a common core machinery. Mol. Cell 46, 542–548.

Zybailov, B., Coleman, M.K., Florens, L., and Washburn, M.P. (2005). Correlation of relative abundance ratios derived from peptide ion chromatograms and spectrum counting for quantitative proteomic analysis using stable isotope labeling. Anal. Chem. 77, 6218–6224.

